# Molecular Basis of Mycoparasitic Performance: Genomic and Transcriptomic Comparison of Contrasting *Trichoderma atroviride* Strains

**DOI:** 10.64898/2026.06.22.733667

**Authors:** Etienne Brémand, Franck Bastide, Justine Colou, Nicolas Denancé, Séverine Boisard, Nicolas Ruiz, Samuel Bertrand, Muriel Marchi, Jérôme Verdier, Thomas Guillemette

## Abstract

*Trichoderma* species are widely used as biological control agents due to their ability to parasitize plant pathogens. However, substantial variability in mycoparasitic performance exists among strains, even within the same species, and the underlying molecular mechanisms remain poorly understood. Here, we performed comparative genomic and transcriptomic analyses of six *Trichoderma atroviride* strains exhibiting contrasting mycoparasitic performance (weakly or highly parasitic; WP or HP) against *Alternaria brassicicola*, *Rhizoctonia solani*, and *Globisporangium ultimum*.

Comparative genomics revealed limited strain-specific differences, mainly restricted to NLR (NOD-like receptor) repertoires, with certain NLR-coding genes absent from WP strain genomes compared to HP strains, while overall genomic variation remained low. In contrast, transcriptomic analyses revealed strong differences in gene expression dynamics between HP and WP strains.

Co-expression network analysis identified two modules associated with mycoparasitic performance. The first was specifically induced in response to pathogen contact and was enriched in genes encoding cell wall-degrading enzymes, with stronger expression in HP strains. The second module was more broadly overexpressed in HP strains across all conditions and included genes involved in detoxification and defense-related pathways. In addition, this module encompassed genes involved in specialized metabolite biosynthesis and effector-like protein secretion, with WP and HP strains differentially expressing distinct gene subsets within these categories.

Together, these results provide a comprehensive framework for identifying the molecular drivers of mycoparasitic performance in *T. atroviride*. This study deepens our understanding of the functional diversity within the species and establishes a robust foundation for the future development of molecular markers to predict strain efficiency.

**Highlights:**

– Significant variability in mycoparasitic performance exists within *Trichoderma atroviride*.
– Certain NLR receptor genes are specific to highly parasitic genomes.
– Highly parasitic strains show stronger expression of CWDE, ROS detoxification, and defense-related pathways.
– Highly and weakly parasitic strains differ in expressed specialized metabolism and effector-like gene sets.

## 1 Introduction

Crop production is challenged by many abiotic constraints and biotic threats. Even before crop establishment, seeds may be affected by pre- or post-emergence damping-off caused by fungi, such as *Alternaria*, *Rhizoctonia* or *Fusarium* species, or by oomycetes including *Globisporangium* and *Phytophthora* species (Lamichhane et al., 2017). Several alternatives to chemical pesticides have been developed to control these pathogens, like the use of mycoparasitic fungi. *Trichoderma* spp. have gained considerable attention as they can act on the pathogen through nutrient competition, antibiosis, and mycoparasitism or on the plant by inducing defense responses (Mukherjee et al., 2013).

The genus *Trichoderma* comprises 400 species distributed across a wide range of ecosystems, including soils, rhizospheres, decaying plant material, living plants, marine habitats and even in immunocompromised humans (Cai and Druzhinina, 2021). Owing to their biocontrol and biotechnological potential, *Trichoderma* species have been the focus of extensive genomic and transcriptomic investigations (Druzhinina et al., 2011), with 239 genome sequences currently available in the NCBI database (in April 2026).

*Trichoderma* core genome comprises approximately 7,000 orthologs, whereas individual strains harbor between 9,000 and 14,000 genes, reflecting substantial genomic plasticity (Kubicek et al., 2019). Among the key components of the *Trichoderma* enzymatic arsenal are glycoside hydrolases (GH), which play a central role in carbohydrate metabolism and fungal cell wall degradation. Several GH families involved in chitin, chitosan, and β-glucan degradation are particularly expanded in *Trichoderma* species compared to other fungi (Druzhinina et al., 2012; Schmoll et al., 2016). In addition, *Trichoderma* species produce a wide array of specialized metabolites, including non-ribosomal peptides, polyketides, terpenes, and volatile organic compounds (Reino et al., 2008), as well as effector-like proteins implicated in interactions with both plants and microbial competitors (Guzmán-Guzmán et al., 2017).

Recently, *Trichoderma atroviride* was shown to be overrepresented in seeds (Brémand et al., 2026). Some strains were able to limit the transmission of seed-borne pathogens to seeds, like *Alternaria brassicicola*, and to reduce the impact of soil-borne pathogens, such as *Globisporangium ultimum*. However, marked variability in biocontrol efficiency was observed among *T. atroviride* strains in that study, both *in vitro* and *in planta*, suggesting strain-specific mechanisms underlying mycoparasitic performance. While transcriptomic studies have shed light on general responses of *T. atroviride* to pathogen contact (Chen et al., 2023; Reithner et al., 2011; Seidl et al., 2009b), little is known about how these mechanisms differ between weakly parasitic (WP) and highly parasitic (HP) strains. Understanding these differences remains a major challenge in optimizing *Trichoderma*-based protective products.

In this study, we investigated six *T. atroviride* strains exhibiting contrasting mycoparasitic performance against *A. brassicicola*, *R. solani*, and *G. ultimum*. We combined genome sequencing with extensive RNA-seq analyses of *in vitro* confrontation assays between the six *T. atroviride* strains and the three pathogens. This integrative genomic and transcriptomic approach provided insights into molecular strategies associated with mycoparasitism and enabled the identification of molecular mechanisms that may contribute to the enhanced performance of highly mycoparasitic strains.

## 2 Materials and Methods

### 2.1 Strains and Cultivation Conditions

*Trichoderma atroviride* strains were obtained from the commercial product TRI-SOIL® (I1237) of Agrauxine by Lesaffre (Beaucouzé, France), isolated from seeds of tomato (P3041), flax (P3080) or lettuce (P3116) by GEVES (Angers, France) and isolated from marine environment (MMS1295 and N1508) by ISOMer (Nantes, France). Three pathogens responsible for damping-off were used in this study. *Alternaria brassicicola* Abra43 (CIRM CF, BRFM3693) isolated from radish seeds by IRHS (Angers, France). *Rhizoctonia solani* PAT-009 isolated from radish by CDDM (Comité Départemental de Développement Maraîcher, Nantes, France). *Globisporangium ultimum* CBS114.19 isolated from gymnosperm seedlings (Westerdijk Fungal Biodiversity Institute). For long-term storage, all strains were kept at −80 °C as mycelial explants stored in 30% glycerol, except *G. ultimum* kept at 4 °C as mycelial explants stored in sterile water. For fresh cultures, all strains were grown on Potato Dextrose Agar (PDA, Biokar, 39 g/L) at 22°C in the dark.

### 2.2 *In Vitro* Confrontation Assays Between *T. atroviride* and Phytopathogenic Agents

A dual culture assay was performed in square Petri dishes (120 mm × 120 mm) containing PDA medium. Plates were inoculated with two 5-mm-diameter mycelial plugs placed at opposite corners of the dish (one from the pathogen and one from *T. atroviride*). Plugs of *A. brassicicola* were placed 3 days prior *T. atroviride*, and simultaneously for *R. solani* and *G. ultimum*. Three replicates were performed for each condition, and the assay was repeated in three independent experiments. Plates were incubated at 20 °C in the dark. The mycoparasitic performance of each *T. atroviride* strain was visually assessed 10 days after contact with the pathogen, using the scale from Brémand et al. (2026). Statistical analyses were performed in R v4.3.1 using a Kruskal-Wallis test followed by pairwise comparisons with Dunn’s post hoc test and Bonferroni correction.

### 2.3 Inhibition of Pathogen Growth by *T. atroviride* Culture Filtrates

Liquid cultures of *T. atroviride* in PDB (Potato Dextrose Broth, Difco, 24 g/L) and their filtration were performed according to Chateau et al. (2024). For *A. brassicicola*, 10^6^ conidia/mL were collected from a 7-day-old culture on V8 agar medium and suspended in sterile water. For *R. solani*, a mycelial suspension was used as inoculum. Mycelium from a 4-day-old culture grown on PDA with a pectocellulosic membrane was collected and blended for 30 s in Phosphate Buffered Saline (PBS) pH 5.5. The suspension was then filtered to retain hyphal fragments between 60 µm and 500 µm. The hyphal suspension was washed three times with PBS. To adjust the concentration, a range of dilutions was prepared, and turbidity was measured in triplicate using a nephelometer (NEPHELOstar, BMG Labtech). The suspension was then adjusted to a defined turbidity value (device-dependent) corresponding to 10⁴ CFU/mL. For *G. ultimum*, 10^4^ oospores/mL were harvested from 15-day old culture on Synthetischer Nährstoffarmer Agar (SNA) medium (Nirenberg, 1976). The sensitivity of the pathogens to *T. atroviride* culture filtrates was tested in liquid medium using 96-well plates and a 635-nm laser nephelometer (NEPHELOstar, BMG Labtech) (Joubert et al., 2010). PDB was inoculated with 10% (v/v) of pathogen suspension and 10% (v/v) of *T. atroviride* culture filtrate in a final volume of 200 μL. Plates were incubated at 25 °C and growth was monitored every 10 min for 35 h for *A. brassicicola* and *G. ultimum*, and 96 hours for *R. solani*. Three replicates were performed for each condition, and the assay was repeated in three independent experiments. The area under the curve (AUC) was calculated for each growth curve. Pairwise comparisons were conducted using Tukey’s Honest Significant Difference (HSD) post hoc test. The percentage of inhibition was determined for each condition containing a *T. atroviride* filtrate relative to the control condition without filtrate.

### 2.4 DNA Sequencing, Genome Annotation, and Comparative Genomics

High-molecular-weight genomic DNA was extracted from 7-day-old cultures of *T. atroviride*, grown on pectocellulosic membranes placed on PDA medium as described by Möller et al. (1992) and adapted by Colou et al. (2026). Genomic DNA was sequenced on a PacBio Revio platform (Novogene, UK). SMRTbell libraries were prepared using standard protocols to generate HiFi reads via Circular Consensus Sequencing. Only HiFi reads with Q30 quality (>99.9% accuracy) were used for *de novo* assembly.

Genome assembly was performed using Flye v2.9.5 2, and assembly quality was assessed with QUAST v5.2.0 (Gurevich et al., 2013) and BUSCO v5.7.1 (Manni et al., 2021) using the “fungi_odb10” database. Genomic similarity among strains was evaluated by computing pairwise distances with Mash v2.3 (Ondov et al., 2016). The resulting distance matrix was used to generate a Newick-formatted tree, which was visualized in MEGA X v10.2.6 (Kumar et al., 2018). The phylogenetic tree includes the six *T. atroviride* strains from this study, ten additional *T. atroviride* strains and one *Trichoderma gamsii* strain (Supplementary Table 1). Gene prediction was performed using Helixer v0.3.4 (Stiehler et al., 2021). The GenBank accession numbers for the assemblies and annotations of the six genomes are provided in Supplementary Table 1. Functional annotation was performed using eggNOG-mapper v2.1.13 (Cantalapiedra et al., 2021) to infer gene functions and identify PFAM (Protein families database) domains (Mistry et al., 2021). Specialized metabolite biosynthetic gene clusters were predicted using fungiSMASH v8.0.4 (Blin et al., 2025). Gene Ontology terms were retrieved through the STRING database v12.0 (Szklarczyk et al., 2025). CAZyme annotation was conducted using dbCAN3 v12 (Zheng et al., 2023). Peptidases were annotated by performing BLASTp searches of the predicted proteome against the MEROPS peptidase database v12.4 (Rawlings et al., 2010). For effector-like protein annotation, secreted proteins were first extracted from the predicted proteome using SignalP v5.0 (Almagro Armenteros et al., 2019). Effector-like proteins were then predicted using EffectorP v3.0 (Sperschneider and Dodds, 2022). The functional annotation of all *T. atroviride* N1508 genes is available in Supplementary Dataset 1. Orthologous groups were inferred between the six *T. atroviride* strains using OrthoFinder v2.5.2 (Emms and Kelly, 2019), with protein alignments performed using Diamond. All orthologue groups, as well as the names of the genes within each group, are available in Supplementary Dataset 2.

### 2.5 RNA Sequencing, Differential Expression, and Co-Expression Network Analyses

*T. atroviride* transcriptomes were analyzed during *in vitro* confrontation assays using the same experimental design as described in section 2.2, but adapted with PDA medium covered with a pectocellulosic membrane. To ensure synchronized contact, the initial spacing between *T. atroviride* and each pathogen was adjusted according to their growth rates and plugs of *A. brassicicola*, *R. solani*, and *G. ultimum* were placed 7 days before, 1 day after, or 2 days after *T. atroviride*, respectively. Self-confrontation of *T. atroviride* (second plug added one day after the first) served as a control. All assays were performed in triplicate. Samples were collected from the colony edge of *T. atroviride* at two time points: before physical contact (four days after *T. atroviride* inoculation) and after contact (seven days after *T. atroviride* inoculation). Fungal tissues were flash-frozen in liquid nitrogen and bead milled. Total RNA was extracted using the NucleoSpin RNA Set for NucleoZOL kit (Macherey-Nagel) and further purified with the RNA Clean & Concentrator-5 kit (Zymo Research). RNA concentration and integrity were measured using a TapeStation system (Agilent Technologies). RNA samples were then sent to Beijing Genomics Institute (BGI, China) for sequencing. cDNA libraries were prepared following BGI’s standard protocols and sequenced using the DNBseq™ technology, generating 20 million paired-end reads (150 bp) per sample.

Reads were aligned to the *T. atroviride* N1508 transcriptome using Salmon v1.10.3 (Patro et al., 2017) to obtain counts and TPM (Transcripts Per Million) values. A multidimensional scaling (MDS) analysis based on Bray-Curtis distances was performed on TPM data in R using the vegan package (v2.6.10). For weighted gene co-expression network analysis (WGCNA), raw RNA-seq counts were transformed using the variance stabilizing transformation from the DESeq2 v1.42.0 package (Love et al., 2014) and genes with variance < 0.3 were removed. Co-expression network modules were identified using the R package WGCNA v1.73 (Langfelder and Horvath, 2008) with the automatic one-step network construction method for module detection. The soft-thresholding power was set to 8, the minimum module size to 30, and the merge cut height to 0.20. A signed hybrid topological overlap matrix was used to calculate gene co-expression similarity. Modules with highly similar eigengenes were merged, and module eigengenes were calculated.

To examine gene-specific expression differences, a classical differential expression analysis was performed using the R package AskoR v1.0.0 (Carvalho et al., 2021). This package relies on edgeR v4.0.3 (Chen et al., 2025) and defines genes as significantly differentially expressed when the absolute log2 fold change > 0.5 and FDR (False Discovery Rate) < 5%. Comparisons were performed between pre-and post-contact conditions using combined datasets from *A. brassicicola*, *R. solani*, and *G. ultimum* for both WP and HP strains. In addition, differential expression analyses were conducted between WP and HP strains using post-contact data from these three pathogens. Supplementary Dataset 1 details each gene presence in a WGCNA expression module as well as the results of the differential expression analysis.

## 3 Results

### 3.1 Phenotype and Mycoparasitic Performance of *T. atroviride* Strains

The *T. atroviride* strains I1237, P3041, P3080, P3116, MMS1295, and N1508 were selected for this study. Strains I1237 and P3041 exhibit thinner and more compact mycelia compared to the other four strains, which develop thicker and more cottony mycelia (Fig. 1A). In confrontation assays, strains I1237 and P3041 exhibited weak parasitic (WP) activity against the three damping-off pathogens tested, contrary to strains P3080, P3116, MMS1295, and N1508 which showed high parasitic (HP) activities (Fig. 1B). On average, mycoparasitic performance was 13%, 0% and 46% for WP strains and 90%, 76% and 99% for HP strains against *Alternaria brassicicola*, *Rhizoctonia solani* and *Globisporangium ultimum*, respectively. Ability of *T. atroviride* to produce antimicrobial compounds deleterious to pathogen growth was tested by nephelometry using culture filtrates from the six strains (Fig. 1C). The two WP strains (I1237, P3041) produced the least effective filtrates against all three pathogens, with an average inhibition of 50%, 66% and 45% for *A. brassicicola*, *R. solani*, and *G. ultimum*, respectively. By contrast, the values for HP strains reached 82%, 91% and 92%, respectively. No significant differences between HP strains and between WP strains were observed.

**Fig. 1:**
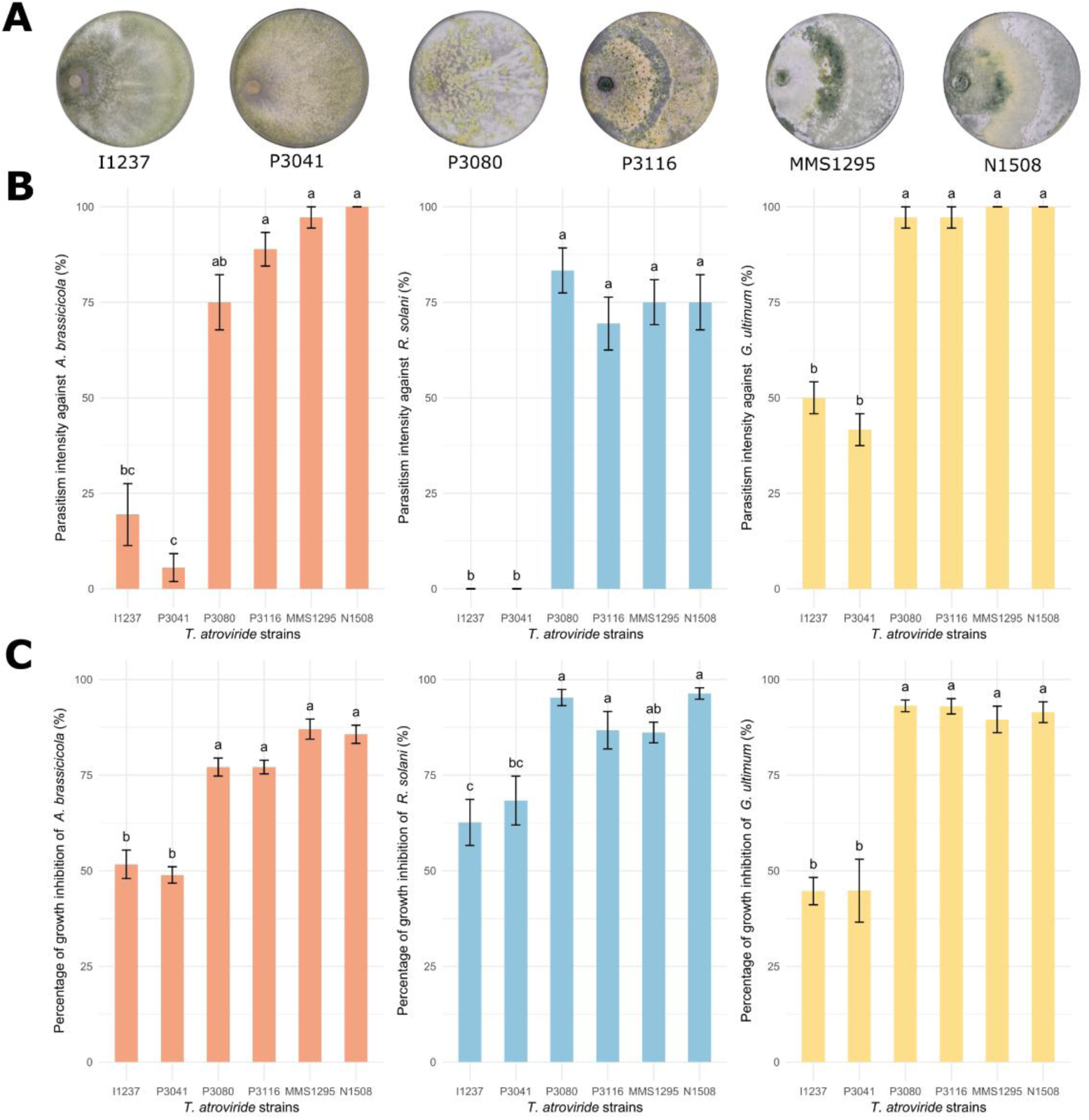
Phenotypic comparison and differences in antagonistic performance among six *Trichoderma atroviride* strains. (A) Photographs of the six strains after 7 days of growth on PDA medium at 22 °C in the dark. (B) Parasitism intensity of the six *T. atroviride* strains against three plant pathogens (*Alternaria brassicicola*, *Rhizoctonia solani* and *Globisporangium ultimum*), determined through dual confrontation assays. (C) Percentage of pathogen growth inhibition caused by culture filtrates from the six *T. atroviride* strains. Error bars represent the standard error, and different letters indicate statistically significant differences (see Materials and Methods for details).

### 3.2 Comparative Genomics of *T. atroviride* Strains

#### 3.2.1 Genome Sequencing, Assembly, and Annotation of Six *T. atroviride* Strains

The genomes of the six *T. atroviride* strains were sequenced. The resulting genome assemblies were of high quality and high completeness (Table 1). A slight difference in genome size was observed among the strains MMS1295 and N1508 having a smaller size (36.7 to 36.9 Mb) compared to the four others (37.2 to 37.5 Mb). Gene annotation identified between 12,440 and 12,491 genes in strains I1237, P3041, MMS1295, and N1508, and slightly more in P3116 (12,530) and P3080 (12,640) (Table 1). The GC content of the genomes ranged from 48.7% to 49.2%. The number of transposable elements varied across the genomes, mostly comprised between 718 and 853, but with P3041 and P3080 appearing as outliers with 354 and 1,033 TE, respectively. This may partly explain why the two strains have the least and the most fragmented genomes, respectively. Approximately 74% of the genes across all strains could be functionally assigned using EggNOG and 73% were annotated with PFAM domains.

**Table 1.** Genome assembly and gene annotation of six *Trichoderma atroviride* strains. Assembly statistics include genome size, number of contigs, N50, mean sequencing coverage, and GC content (%). Transposable element (TE) number and genome proportion (TE coverage) were estimated for each strain with EDTA. Genome completeness was assessed using BUSCO single-copy genes. Predicted protein-coding genes were annotated with EggNOG to obtain gene functions and PFAM domains.

|  | I1237 | P3041 | P3080 | P3116 | MMS1295 | N1508 |
| --- | --- | --- | --- | --- | --- | --- |
| Assembly size (Mb) | 37.2 | 37.2 | 37.5 | 37.5 | 36.7 | 36.9 |
| Number of contigs | 11 | 9 | 52 | 33 | 22 | 16 |
| N50 (Mb) | 5.6 | 5.6 | 2.0 | 3.7 | 3.1 | 3.4 |
| Mean coverage | 35 | 37 | 54 | 77 | 87 | 61 |
| GC content (%) | 48.7 | 48.7 | 48.9 | 48.7 | 49.2 | 49.0 |
| TE number | 847 | 354 | 1,033 | 853 | 814 | 718 |
| TE coverage (%) | 2.3 | 1.4 | 2.5 | 2.2 | 2.3 | 1.6 |
| Busco single-copy (%) | 99.1 | 99.1 | 99.2 | 99.3 | 99.1 | 99.1 |
| Number of predicted genes | 12,44 | 12,445 | 12,64 | 12,53 | 12,486 | 12,491 |
| Genes with EggNOG fonction | 9,16 | 9,171 | 9,246 | 9,194 | 9,193 | 9,201 |
| Genes with PFAM domain | 9,065 | 9,077 | 9,155 | 9,103 | 9,095 | 9,1 |

A genomic similarity tree constructed with Mash confirmed that all six strains belong to the *T. atroviride* species (Fig. 2). The four strains I1237, P3041, P3080, and P3116 are highly similar to each other, while MMS1295 and N1508 form a distinct group that is closely related to each other.

**Fig. 2.**
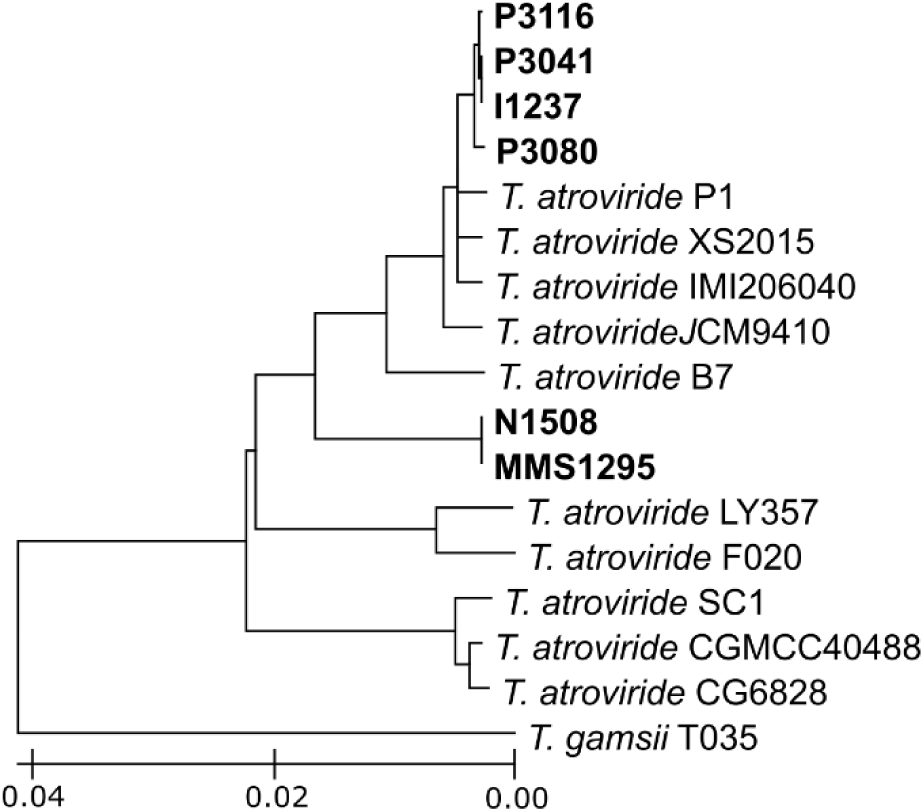
Genomic similarity tree of the six *Trichoderma atroviride* strains compare with references. Genomic similarity tree based on Mash distances calculated from the complete genomes of our six *T. atroviride* strains (in bold) and 10 reference strains of *T. atroviride* (Supplementary Table 1). The tree was generated from the Mash distance matrix, with *Trichoderma gamsii* T035 as the outgroup. The scale represents the Mash distance.

#### 3.2.2 Comparison of Orthologous Groups Between *T. atroviride* Genomes

A total of 13,872 orthologous groups were identified, of which 76% contained at least one gene from each strain (Fig. 3). Differences in orthologous group composition reflect phylogenetic relationships: MMS1295 and N1508 share 82% of orthologous groups, whereas I1237, P3041, P3080, and P3116 share 80%. Functional annotation reflects gene conservation, as 79% of the 10,591 orthologous groups shared by all six strains had an assigned function based on eggNOG annotation, whereas only 15% of the remaining 3,281 orthologous groups were functionally annotated.

**Fig. 3.**
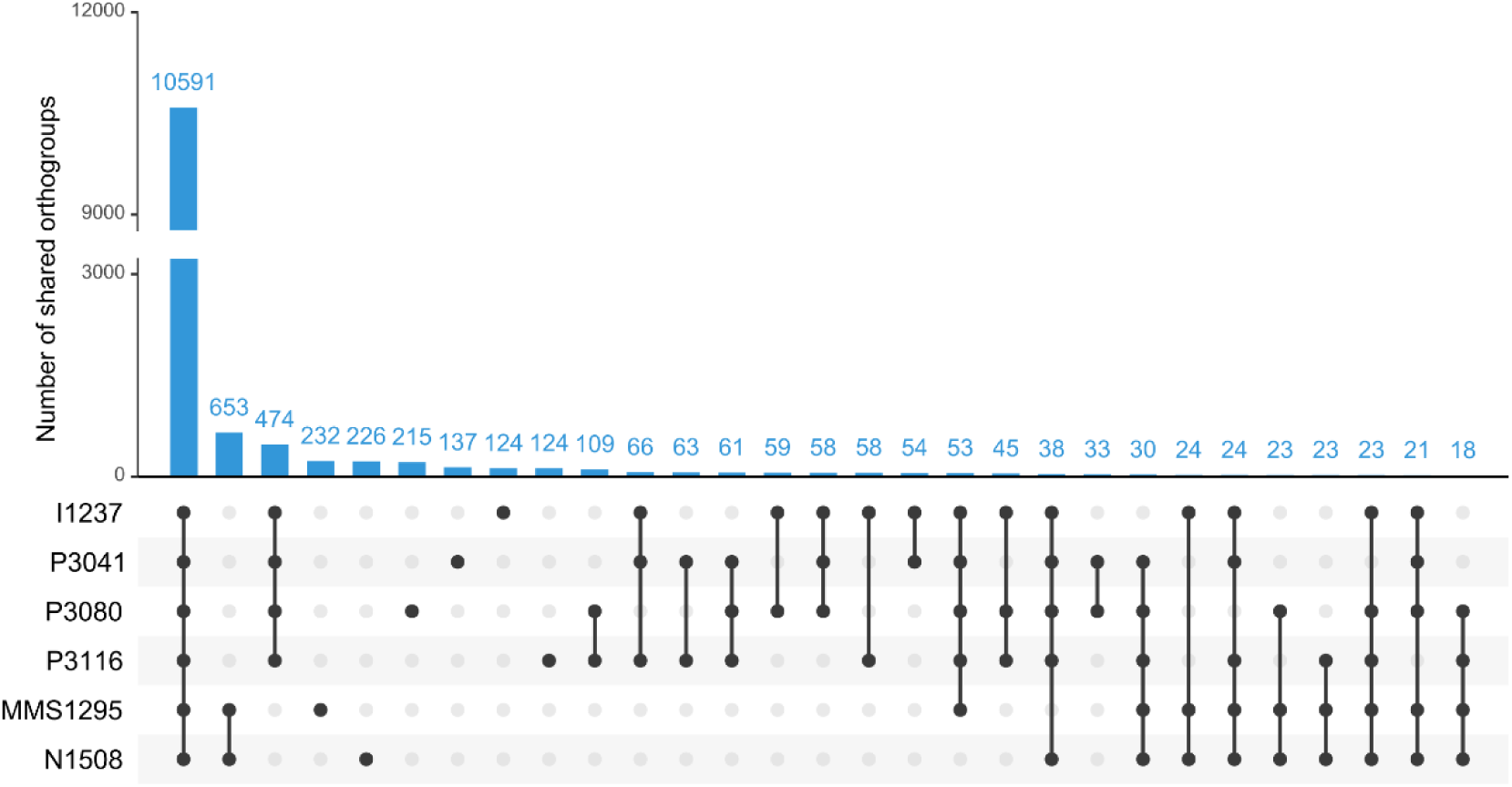
Upset plot of orthologous groups shared among the six *Trichoderma atroviride* strains. An orthogroup was considered shared if at least one gene from each strain was present in that orthogroup. Only the 23 conditions with the highest number of shared orthogroups are shown.

The analysis identified 18 orthologous groups exclusive to the four HP strains, of which only two had functional annotations. These two proteins contain N-terminal domains associated with NACHT ATPases, a feature commonly found in fungal NLR (NOD-like receptor) proteins involved in allorecognition and defense-related signaling (Bonometti et al., 2025; Dyrka et al., 2014). In addition, two orthologous groups were found to be expanded in HP compared with WP strains. The first contained 13 to 14 paralogs in HP strains compared with 12 in WP strains and was generally characterized by the presence of NACHT domains, ankyrin repeats, and occasionally PNP_UDP domains. The second contained 3 to 7 paralogs in HP strains, compared with only two copies in WP strains, and was consistently associated with NACHT domains and WD40 repeats. Notably, only these two orthologous groups showed expansion in HP strains compared with WP strains, and both displayed domain architectures characteristic of fungal NLR proteins.

### 3.3 Transcriptional Responses of *T. atroviride* During Confrontation

#### 3.3.1 RNA Sequencing of *In Vitro* Confrontation Assays

RNA-seq analyses were performed directly on confrontation assays between the six *T. atroviride* strains and the three pathogens (*A. brassicicola*, *R. solani*, and *G. ultimum*) or a self-confrontation of each *T. atroviride* strain. Samples were collected before contact, when the two fungi were 1–2 cm apart, and after contact, when the HP *T. atroviride* strains began to overgrow the pathogen. Interestingly, RNA extracted from samples collected following contact between a HP *T. atroviride* strain and one of the three pathogens was generally more degraded than in other conditions (Supplementary Fig. 1). Reads were subsequently mapped to the transcriptome of either *T. atroviride* N1508 or the corresponding pathogen. The N1508 genome was chosen because it is the least fragmented among the HP strains. On average, 91% and 66% of the reads were aligned to N1508 transcriptome in samples collected before and after pathogen contact, respectively (Supplementary Fig. 2). This drop could be explained by a fraction of reads mapping to the pathogen transcriptome, but mainly by reads that could not be mapped to either transcriptome. In line with phylogenetic relationships, the proportion of reads mapping to both *T. atroviride* and the pathogen under post-contact conditions averaged 3% for *A. brassicicola* (ascomycete), 0.036% for *R. solani* (basidiomycete), and 0.008% for *G. ultimum* (oomycete). Those reads were removed from the analysis. Multidimensional Scaling (MDS) of the samples grouped, on the first axis, strains two-by-two: the WP strains I1237/P3041, the seed-borne HP strains P3080/P3116 and the marine HP strains MMS1295/N1508 (Fig. 4). This is consistent with their phylogenetic proximity and their mycoparasitic performance. The second axis primarily captures differences between pre- and post-contact samples, located at the top and bottom, respectively.

**Fig. 4.**
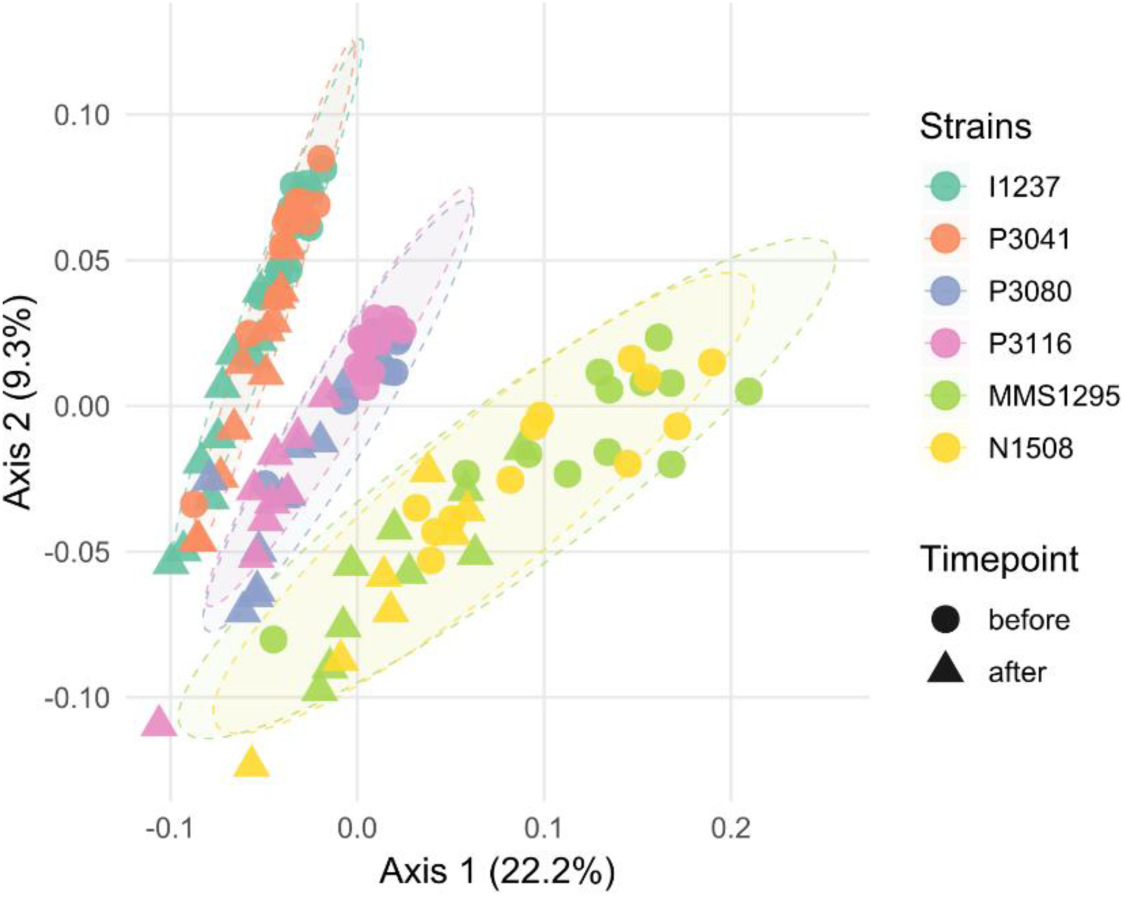
MDS of RNA-Seq samples based on TPM-normalized gene expression. Colors indicate different *Trichoderma atroviride* strains. Circles represent samples before contact, and triangles represent samples after contact with pathogen. Ellipses represent the 95% confidence interval around the samples of each strain.

#### 3.3.2 Co-Expression Network Analysis

A weighted gene co-expression network analysis (WGCNA) was performed to group co-expressed genes, defined as genes displaying similar expression patterns across samples, into expression modules based solely on correlation structure. Of the 12,491 genes identified in the *T. atroviride* N1508 transcriptome, 54% exhibited sufficient expression variability across samples and were retained for analysis, resulting in their assignment to 30 distinct expression modules. These modules were further grouped into seven higher-order expression modules (EM) based on the similarity of their expression profiles (Supplementary Fig. 3). Functional enrichment analysis based on Gene Ontology (GO) terms was performed to identify biological processes associated with each expression module (Supplementary Fig. 4-10). The co-expression network (Fig. 5A) displays individual modules as distinct colors, organized into these seven major EM.

**Fig. 5.**
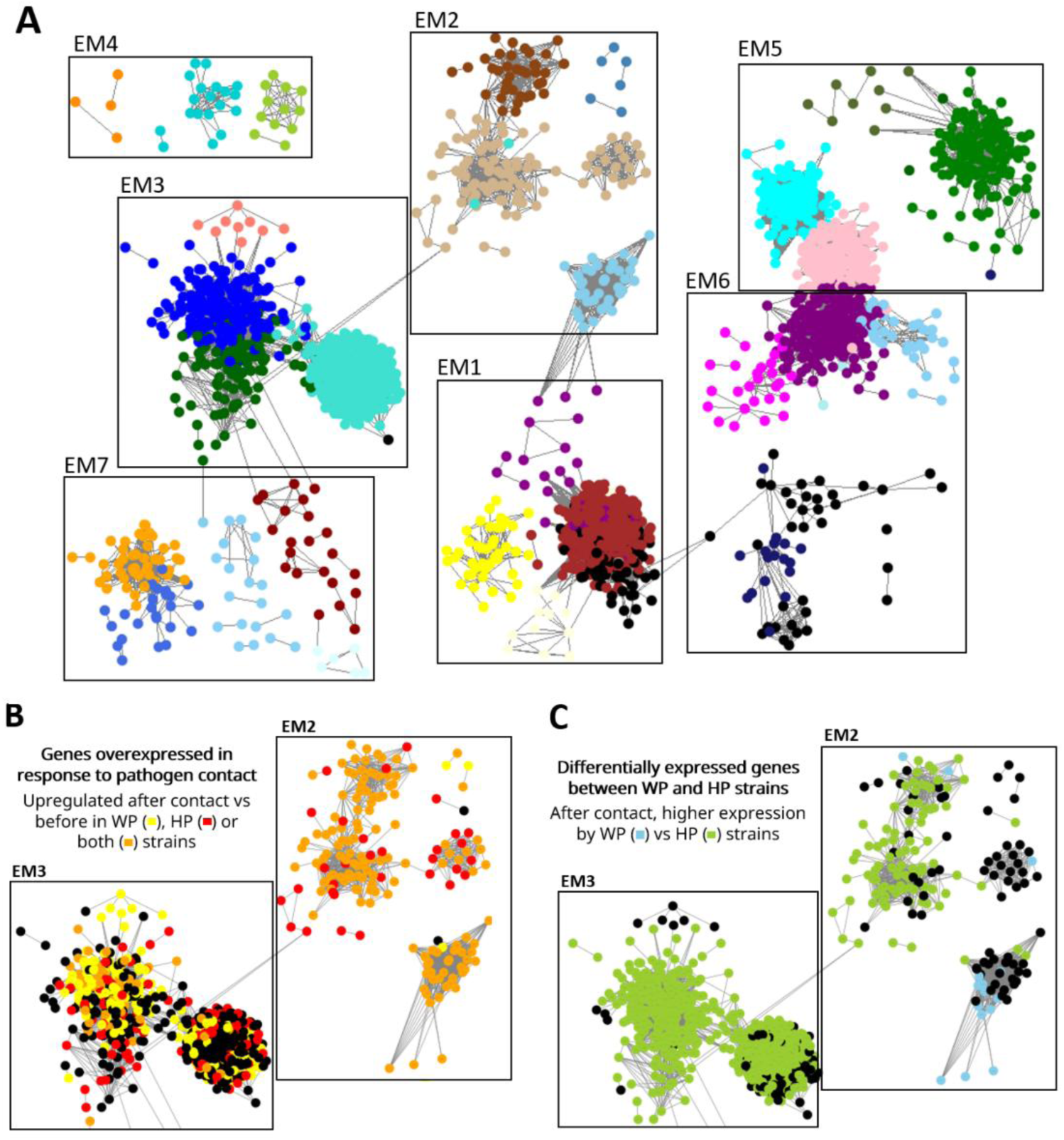
Co-expression network derived from WGCNA analysis. The network was visualized using Cytoscape, considering only correlations above 0.15. At the top, node colors indicate WGCNA modules, which were further grouped into seven major expression modules (EM) based on similarity in expression profiles (A). At the bottom, node colors represent results from differential expression analyses focusing on EM 2 and 3. On the left, genes upregulated following pathogen contact compared to pre-contact conditions are shown (B). On the right, genes differentially expressed after pathogen contact between weakly parasitic (WP) and highly parasitic (HP) strains are displayed (C). In black, non-differentially expressed genes.

Figure 5B displays, for EM2 and EM3, the genes significantly differentially expressed between before and after pathogen contact. EM2 is strongly associated with pathogen contact in both HP and WP strains. Differential expression analysis further indicates that a subset of genes in EM3 is associated with pathogen contact in both strain types, whereas the response observed in EM1 is predominantly restricted to WP strains (Supplementary Fig. 4A). Figure 5C shows, after pathogen contact, expression differences between WP and HP strains. EM3 is primarily composed of genes more highly expressed in HP strains. Similarly, EM2, which is strongly associated with pathogen contact, also contains a substantial proportion of genes upregulated in HP strains. In contrast, EM1, which is induced in response to pathogen contact in WP strains, is also more highly expressed in WP strains compared to HP strains (Supplementary Fig. 4B). A similar trend is observed for EM6, which is also more highly expressed in WP strains.

EM2, which is predominantly induced upon pathogen contact and exhibits higher expression in HP strains, comprises 451 genes. Gene Ontology (GO) enrichment analysis revealed that this EM is enriched in biological processes related to lipid and nitrogen compound catabolism, as well as carbohydrate degradation, including β-glucan, chitin, cellulose, hemicellulose and xylan breakdown, and more broadly cell wall degradation processes (Supplementary Fig. 5). EM3, which is globally more highly expressed in HP strains compared to WP strains, contains 1,593 genes. Enriched GO terms in this module are associated with carbohydrate metabolism (including monosaccharide, pentose and arabinose catabolism), specialized metabolite biosynthesis, and detoxification processes, such as responses to reactive oxygen species (ROS) and toxic compounds (Supplementary Fig. 6).

Based on these observations, subsequent analyses focused on the functional categories enriched in these two EM, examining both their genomic distribution across strains and their expression dynamics across conditions (before *versus* after pathogen contact) and between strain types (HP *versus* WP).

### 3.4 Molecular Mechanisms Involved in Mycoparasitic Performance

#### 3.4.1 Cell Wall-Degrading Enzymes

Enzymes involved in carbohydrate degradation are likely to contribute to host cell wall breakdown, enabling *T. atroviride* to kill its fungal host and subsequently exploit it as a nutrient source. To further investigate this potential, carbohydrate-active enzymes (CAZymes) were systematically identified. Across the six *T. atroviride* strains investigated, 256-258 glycoside hydrolases (GH), 100 glycosyltransferases (GT), 10 polysaccharide lyases (PL), 24 carbohydrate esterases (CE), and 72 auxiliary activity enzymes (AA) were detected (Fig. 6A). Transcriptomic analyses revealed a strong association between CAZyme expression and pathogen interaction, with 67% of CAZyme-encoding genes being overexpressed in response to pathogen contact in WP or HP strains (Fig. 6A). Among all these genes, 32% were more highly expressed in HP than in WP strains after pathogen contact, whereas 24% were more highly expressed in WP strains. GH showed an even stronger response to pathogen interaction, with 77% of genes being overexpressed in response to pathogen contact in either WP or HP strains. AA, CE, and PL also seem to be strongly associated with pathogen contact compared with GT.

**Fig. 6.**
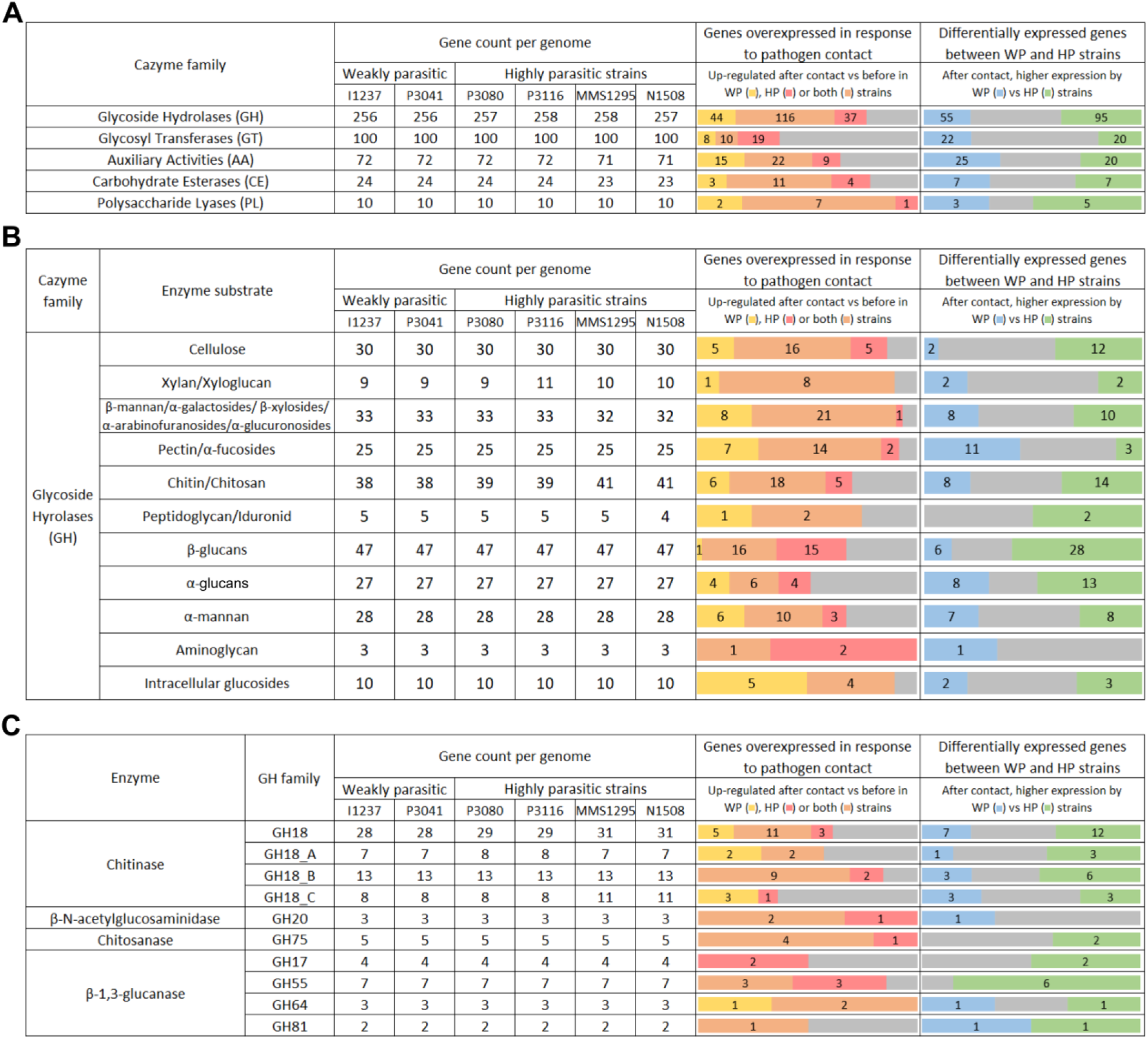
Number of CAZymes genes in each genome and their expression. The central panel shows the number of genes identified in each of the six *Trichoderma atroviride* genomes. On the right, the number of genes overexpressed in post-contact conditions with a pathogen compared to pre-contact conditions is indicated for weakly parasitic (WP), highly parasitic (HP) strains, or both. The far-right panel shows the number of genes more highly expressed by WP or HP strains under post-contact conditions with a pathogen. In grey, non-differentially expressed genes. RNA-seq reads were mapped to the N1508 transcriptome, so gene counts in the expression analysis correspond to N1508 genes. Genes are grouped by CAZyme family (A), as well as by substrate for glycoside hydrolases (B) (details in Supplementary Fig. 13). Enzymes involved in fungal cell wall degradation are further detailed (C). For GH18, the three groups (A, B, and C) were identified by sequence comparison with references (Supplementary Fig. 14).

GH were further classified according to their substrate specificity (Fig. 6B; Supplementary Fig. 13). Genes overexpressed in response to pathogen contact were distributed across all functional groups, being related to fungi or oomycete cell wall degradation or not. Higher expression of GH in HP compared to WP strains is a hallmark of enzymes involved in β-glucan (60%), chitin/chitosan (34%), and cellulose (40%) degradation. GH18 chitinases, the most abundant family in *T. atroviride* genomes are present in variable numbers in I1237/P3041 (28 genes), P3080/P3116 (29), and MMS1295/N1508 (31) (Fig. 6C). Overall, these chitinases are distributed across 23 orthologous groups. The additional GH18 genes in P3080/P3116 results from the expansion of a single orthologous group, which contains three paralogues instead of two, while the extra ones in MMS1295/N1508 belong to three distinct orthologous groups specific to these strains.

GH18 chitinases are classified into three major groups (A, B, and C) based on sequence features, each further subdivided into subgroups according to conserved motifs in their catalytic sites (Seidl et al., 2005) (Supplementary Fig. 12). The *T. atroviride* strains count members of groups A (7-8 genes), B (13), and C (8-11) (Fig. 6C). Among group A chitinases, two genes are consistently overexpressed in response to pathogen contact across all strains, while two additional genes are exclusively in WP strains. Following contact, three group A chitinases show higher expression in HP strains, compared to one in WP strains. In P3080/P3116, the additional GH18 is an A5-type chitinase, this gene could not be included in the differential expression analysis, as RNA-seq reads were mapped to the N1508 reference transcriptome which does not contain it. Group B chitinases are the most abundant (13 members) and exhibit the strongest association with pathogen contact, with 11 overexpressed genes. Among them, six are more highly expressed in HP strains and three in WP strains, respectively. In contrast, group C chitinases appear less responsive to pathogen interaction, with only three genes overexpressed in response to pathogen contact for WP strains and only one for HP strains. Nevertheless, differences between strains exist: three group C chitinases are more highly expressed in HP strains after pathogen contact, while three others are more highly expressed in WP strains. In MMS1295 and N1508, the three additional chitinases are one C1-type with uniform expression under the conditions tested, and two C2-type with higher expression in HP compared to WP strains but without induction in responses to pathogen contact.

The six genomes contain β-1,3-glucanases from four families: four GH17, seven GH55, three GH64, and two GH81 enzymes (Fig. 6C). Their expression profiles differ markedly between strains. Of them, several are overexpressed in response to pathogen contact: six in all strains, one (GH64) only in WP strains, and five (three GH55, two GH17) exclusively in HP strains. When comparing HP and WP strains after pathogen contact, ten β-1,3-glucanases (including five GH55) show higher expression in HP strains compared to two only (GH64, GH81) in WP strains. Notably, no GH17 genes are induced in response to pathogen contact in WP strains. Genes encoding β-1,3-glucanases show the greatest variation among strains, with a substantial proportion being more highly expressed in HP strains.

Peptidases may contribute to mycoparasitism by degrading protein components of the host cell wall, thereby facilitating the action of polysaccharide-degrading enzymes. The MMS1295 and N1508 strains encoded the lowest numbers of peptidases (433 and 434, respectively) compared to the others (447 or 448) (Fig. 7). However, secreted peptidases, that account for 26-28% of the total repertoire, are more abundant in MMS1295 and N1508 (120 and 121, respectively *versus* 114 to 116). Overall, 62% of secreted peptidase genes were overexpressed in response to pathogen contact in WP or HP strains, and 43% showed higher expression in HP compared to WP strains following pathogen contact. Secreted peptidases can be distinguished into catalytic classes based on the nature of their active site. The six genomes predominantly contain serine peptidases (66–75 genes), aspartic peptidases (18–19) and metallopeptidases (17–18) (Fig. 7). A relatively consistent proportion of these enzymes was induced upon pathogen contact. Notably, subtilases (S08) were strongly overexpressed in response to pathogen contact, with 11 of the 14 genes being upregulated in either WP or HP strains, and 6 showing higher expression in HP than in WP strains following pathogen exposure.

**Fig. 7.**
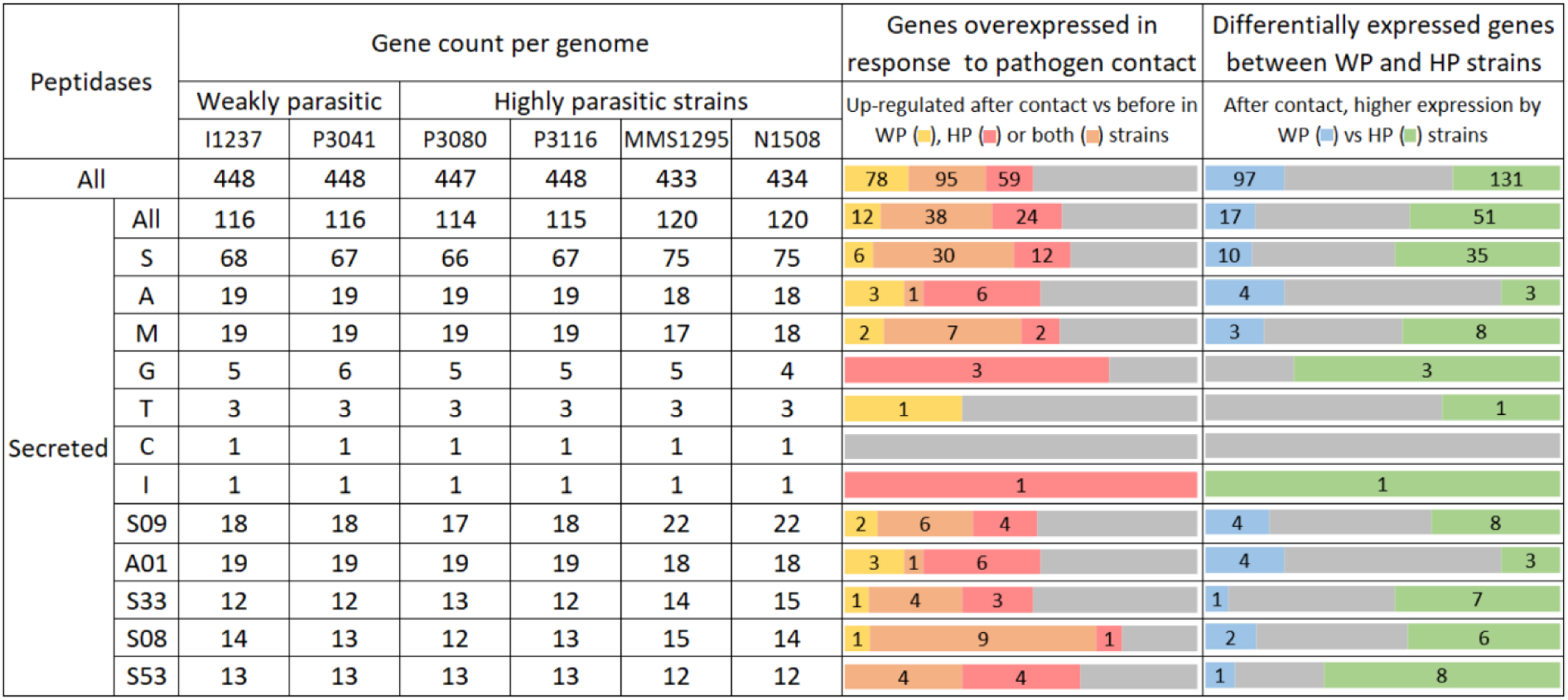
Number of peptidase genes in each genome and their expression. The central panel shows the number of genes identified in each of the six *Trichoderma atroviride* genomes. On the right, the number of genes overexpressed in post-contact conditions with a pathogen compared to pre-contact conditions is indicated for weakly parasitic (WP), highly parasitic (HP) strains, or both. The far-right panel shows the number of genes more highly expressed by WP or HP strains under post-contact conditions with a pathogen. In grey, non-differentially expressed genes. RNA-seq reads were mapped to the N1508 transcriptome, so gene counts in the expression analysis correspond to N1508 genes. Peptidases are classified into catalytic classes: aspartic (A), cysteine (C), metallo- (M), serine (S), glutamic (G), threonine (T), asparagine (N), mixed catalytic (P), and inactive homologs (I). They are further subdivided into families, the five most abundant families in secreted peptidases are shown.

Overall, the expression of a large proportion of CAZyme and peptidase genes identified in *T. atroviride* genomes is associated with pathogen contact, underscoring their central role during mycoparasitism. Moreover, the higher expression of many of these genes in HP compared to WP strains following pathogen contact may contribute to their enhanced mycoparasitic performance.

#### 3.4.2 Specialized Metabolites Biosynthesis

Using fungiSMASH, 46 to 48 specialized metabolite biosynthetic gene clusters (BGCs) were identified per *T. atroviride* genome, with terpene synthase (11-12 TPS), polyketide synthases (11 PKS), and non-ribosomal peptide synthetases (7-8 NRPS and 7 NRPS-like) being the most abundant BGC (Supplementary Table 2). Notable variation was observed among the genomes, including both gene-level differences (*e.g.*, the presence of an NPR-metallophore gene only in P3116 and MMS1295) and cluster-level differences (*e.g.*, the absence of an isocyanide biosynthetic cluster in MMS1295 and N1508). When considering the number of biosynthetic genes, differences were also observed among strains, with 57 to 61 genes involved in specialized metabolite biosynthesis (Fig. 8). Among them, 70% were overexpressed in response to pathogen contact in WP or HP strains. These genes predominantly encode PKS, NRPS, and NRPS-like enzymes. Following pathogen contact, 32% and 40% of these genes showed higher expression in HP and WP strains, respectively. These results suggest that the HP strains’ performance may be due to a metabolic profile distinct from WP, but not necessarily to the production of a greater number of specialized metabolites.

**Fig. 8.**
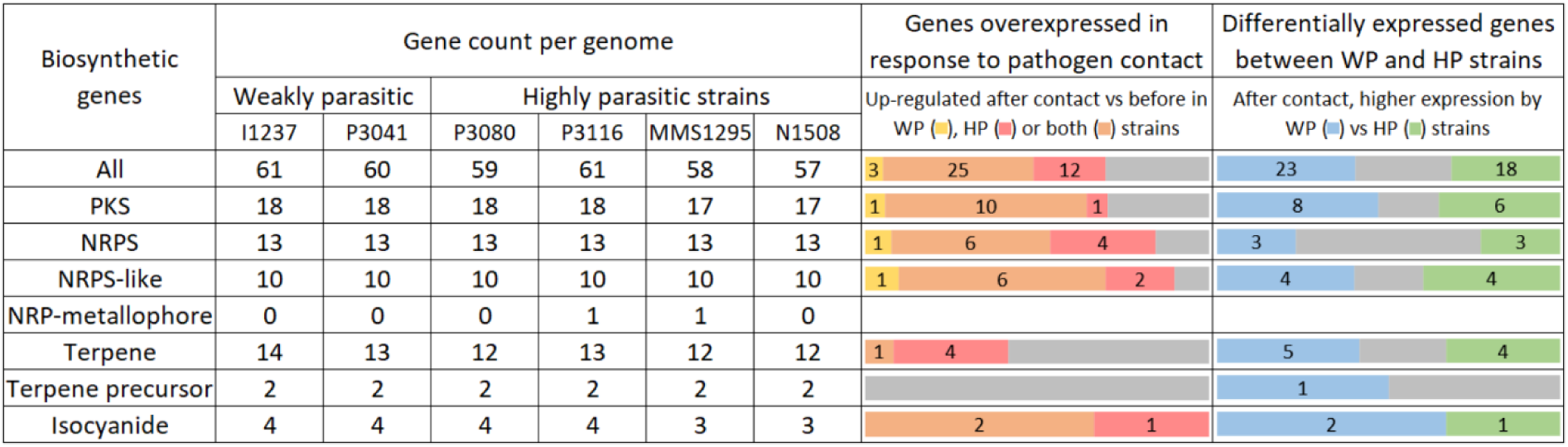
Number of biosynthetic genes in each genome and their expression. The central panel shows the number of genes identified in each of the six *Trichoderma atroviride* genomes. On the right, the number of genes overexpressed in post-contact conditions with a pathogen compared to pre-contact conditions is indicated for weakly parasitic (WP), highly parasitic (HP) strains, or both. The far-right panel shows the number of genes more highly expressed by WP or HP strains under post-contact conditions with a pathogen. In grey, non-differentially expressed genes. RNA-seq reads were mapped to the N1508 transcriptome, so gene counts in the expression analysis correspond to N1508 genes. PKS = Polyketide Synthase, NRPS = Non-Ribosomal Peptide Synthetase.

Notably, in one BGC comprising 26 genes, 14 were more highly expressed in HP compared to WP strains after pathogen contact. They include 4 core biosynthetic genes (two PKS, one NRPS, and one NRPS-like), 5 genes annotated as “biosynthetic-additional”, one Zn(II)₂Cys₆ transcription factor, and other genes potentially involved in biosynthesis (including an aminotransferase and a nitronate monooxygenase). Furthermore, the four core biosynthetic genes were also overexpressed in response to pathogen contact. Additionally, the *pks1* gene, involved in 6-pentyl-α-pyrone (6-PP) biosynthesis, was upregulated in response to pathogen contact. This gene also exhibited higher expression levels in HP than in WP strains following pathogen contact. This polyketide is a volatile compound characterized by a distinct coconut-like odor and is well known for its antimicrobial properties (Flatschacher et al., 2025).

#### 3.4.3 Detoxification Enzymes and Stress Response Pathways

Proteins carrying annotations associated with detoxification enzymes and stress response pathways were identified. Regarding ROS detoxification, each genome contains six superoxide dismutase, 11 peroxidases, one peroxiredoxin and seven (or nine in MMS1295 and N1508) catalases (Fig. 9). Among these 27 genes, 14 were overexpressed in response to pathogen contact in HP or WP strains. When comparing HP and WP strains after pathogen contact, 14 genes were more highly expressed in HP strains, compared to seven in WP strains. Enzymes primarily involved in redox homeostasis were also partially upregulated in response to pathogen contact but exhibited less variation in expression between HP and WP strains.

**Fig. 9.**
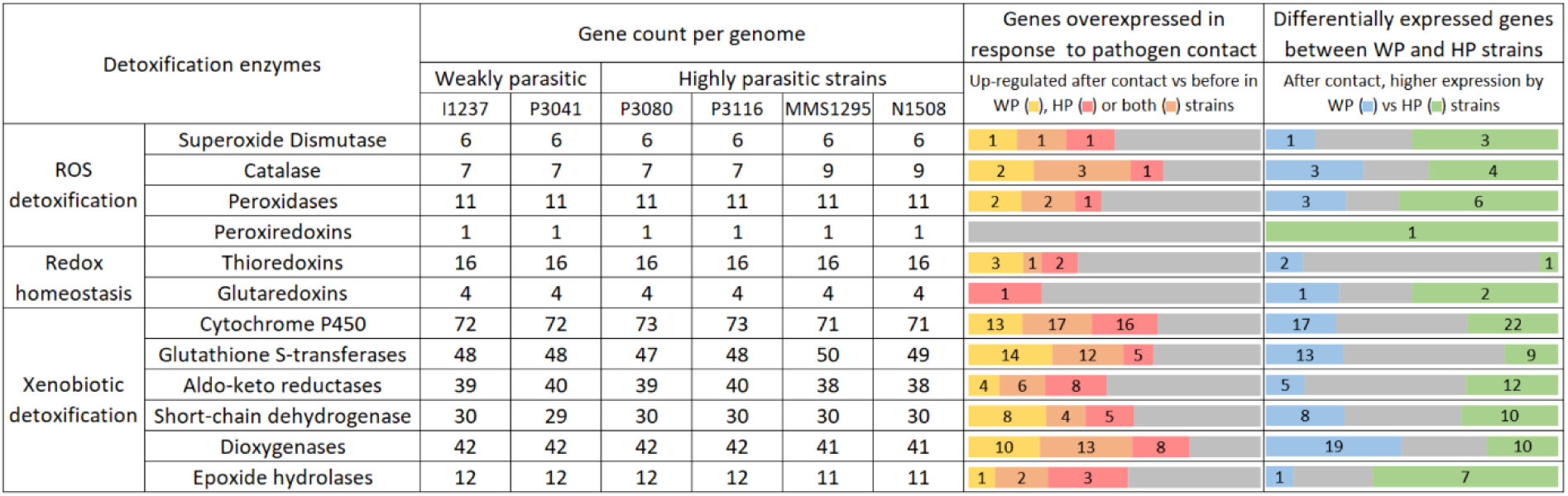
Number of genes involved in detoxification in each genome and their expression. The central panel shows the number of genes identified in each of the six *Trichoderma atroviride* genomes. On the right, the number of genes overexpressed in post-contact conditions with a pathogen compared to pre-contact conditions is indicated for weakly parasitic (WP), highly parasitic (HP) strains, or both. The far-right panel shows the number of genes more highly expressed by WP or HP strains under post-contact conditions with a pathogen. In grey, non-differentially expressed genes. RNA-seq reads were mapped to the N1508 transcriptome, so gene counts in the expression analysis correspond to N1508 genes.

Genes involved in xenobiotic detoxification were predominantly upregulated following exposure to the pathogens, with 62% of the identified genes showing increased expression in WP or HP strains (Fig. 9). Overall, post-contact expression patterns exhibited limited variation between WP and HP strains, particularly for cytochrome P450s, glutathione S-transferases, aldo-keto reductases, and short-chain dehydrogenases. In contrast, a substantial proportion of dioxygenases appeared to be more highly expressed in WP strains, whereas most epoxide hydrolases showed higher expression in HP strains. Taken together, these results suggest that xenobiotic detoxification enzymes are involved in the interaction with the pathogens but display less variation in expression between strains compared to ROS detoxification-related enzymes.

Peroxisomes are essential for reactive oxygen species detoxification and lipid metabolism. Peroxin genes (PEX), involved in their biogenesis, maintenance, and function, are differentially expressed between our conditions (Supplementary Fig. 13). Among the 13 PEX genes identified in the genome, seven were overexpressed in response to pathogen contact in HP strains (PEX2, PEX3, PEX5, PEX11, PEX12, PEX13, and PEX14), compared to only one in WP strains (PEX11). In addition, six genes (PEX1, PEX3, PEX7, PEX11, PEX12, and PEX16) were significantly upregulated in HP strains compared to WP strains after pathogen contact. Other peroxisome-associated enzymes were also more highly expressed in HP strains, including genes involved in β-oxidation (AMACR and SCPx), detoxification (superoxide dismutases, catalases and epoxide hydrolase), and Woronin body formation (HEX1).

Several components of the High Osmolarity Glycerol (HOG) signaling pathway were more highly expressed after pathogen contact in HP strains compared to WP strains, suggesting an enhanced activation of osmotic and oxidative stress responses (Supplementary Fig. 14). This increase mainly involved the Sln1-dependent branch, with higher expression of the response regulator Ssk1 gene, the phosphotransfer protein Ypd1 gene, and the MAPK HOG1 gene. Several downstream transcription factors, including Sko1 and Mcm1, were also more highly expressed in HP strains, as were stress-response effectors such as Gre2 (redox/detoxification) and Cat1 (hydrogen peroxide degradation). Together, these results indicate a coordinated upregulation of the Sln1-Hog1 pathway in HP strains.

Other stress-related pathways were also investigated. In the Cell Wall Integrity (CWI) pathway, upstream components such as PKC1 and RHO1 were not differentially expressed between WP and HP strains. However, several downstream genes, including FKS2, SWI6, MIH1, and CDC28, were more highly expressed in HP strains (Supplementary Fig. 14). Similarly, in the calcineurin pathway, no differential expression was observed for core components (CNA1 and CNB1), whereas the downstream transcription factor CRZ1 was overexpressed in HP compared to WP strains.

Overall, these results indicate that ROS detoxification and stress-response mechanisms are more strongly activated in HP strains. This enhanced stress tolerance likely contributes to their improved mycoparasitic performance.

#### 3.4.4 Effector-Like Proteins

The predicted secretome of the six *T. atroviride* strains contains between 1,072 and 1,093 proteins per genome (Fig. 10). In response to pathogen contact, 54% of the corresponding genes were upregulated in WP or HP strains, and 33% showed higher expression in HP compared to WP strains after pathogen contact.

**Fig. 10.**
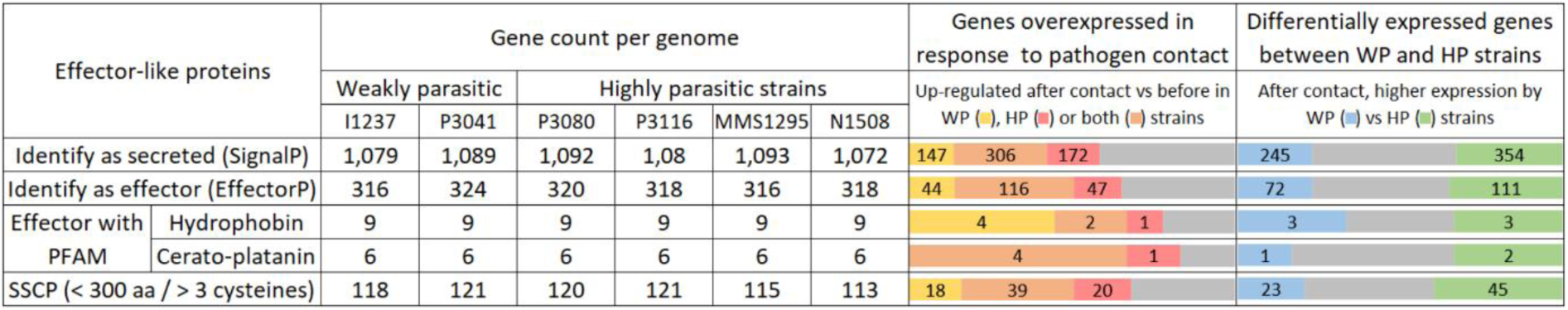
Number of predicted effector-like proteins in each genome and their expression. The left side shows the number of proteins predicted to be secreted by SignalP, proteins predicted as effectors by EffectorP, effectors containing a known PFAM domain, and small secreted cysteine-rich proteins (SSCP: < 300 amino acids, > 3 cysteines and no recognizable PFAM domain) in the genomes of the six *Trichoderma atroviride* strains. The central panel shows the number of genes identified in each of the six *T. atroviride* genomes. On the right, the number of genes overexpressed in post-contact conditions with a pathogen compared to pre-contact conditions is indicated for weakly parasitic (WP), highly parasitic (HP) strains, or both. The far-right panel shows the number of genes more highly expressed by WP or HP strains under post-contact conditions with a pathogen. In grey, non-differentially expressed genes. RNA-seq reads were mapped to the N1508 transcriptome, so gene counts in the expression analysis correspond to N1508 genes.

Using EffectorP, 316-324 effector-like protein candidates per strain were identified. As in the complete secretome, 56% of the effector-like genes were upregulated in response to pathogen contact and 35% showed higher expression in HP strains, compared to 23% in WP strains. Effector-like proteins exhibited high variability in expression profiles across strains, with approximately 60% of these genes being differentially expressed between WP and HP strains. Among these potential effectors, some proteins contain domains commonly associated with fungal effectors, such as hydrophobin and cerato-platanin domains, and most proteins carrying these domains are upregulated in response to pathogen contact (Fig. 10).

Most predicted effector-like proteins lack recognizable PFAM domains. Based on their size (< 300 amino acids) and cysteine content (> 3 cysteines), these proteins were classified as small, secreted cysteine-rich proteins (SSCP). The six genomes contain between 113 and 121 SSCPs (Fig. 10). Among the 113 SSCP identified in strain N1508, 62 were upregulated in response to pathogen contact in WP or HP strains. Forty percent of these SSCP were more highly expressed in HP strains, while 20% were more highly expressed in WP strains following pathogen contact. The 22 SSCP overexpressed exclusively in HP strains in response to pathogen contact, and also more highly expressed in HP than in WP strains following contact, represent good candidates for functional validation.

Overall, a substantial proportion of the identified effector-like proteins appear to be induced in response to pathogen contact. However, these results suggest that the repertoire of expressed effector-like proteins differs between WP and HP strains.

## 4 Discussion

This study shows strong variation in mycoparasitic performance within a single *Trichoderma* species. In a previous study, we reported interspecific variability across several *Trichoderma* species, as well as intraspecific variation within *T. atroviride*, *T. gamsii*, *T. longibrachiatum*, and *T. bissettii* (Brémand et al., 2026). Our results are consistent with their intraspecific findings, as strains P3041, P3080, and P3116 show the same ranking of mycoparasitic performance. In addition, *T. atroviride* strains I1237 (syn. C04) and MMS1295 (syn. M03) were tested in our study and showed patterns consistent with previous reports against *Botrytis cinerea* (Chateau et al., 2024) and *Fusarium* spp. (Pascouau et al., 2023). Moreover, in that same study, the inhibition of *B. cinerea* growth by culture filtrates from strains I1237 and MMS1295 yielded results consistent with our findings. However, it was also shown that the efficiency of culture filtrates can vary markedly depending on culture conditions: by modifying temperature and nutrient availability, the I1237 filtrate could be enhanced to match the efficacy of MMS1295, or conversely, the MMS1295 filtrate could become ineffective. These results suggest that strains with low baseline efficacy are not intrinsically limited in their biocontrol potential, but rather possess the genetic capacity for high antagonistic activity that may be differentially expressed.

Consistent with this interpretation, phylogenomic analyses revealed that the WP strains (I1237 and P3041) are genetically very close to other strains (P3080 and P3116) that exhibit markedly higher mycoparasitic performance. A total of 13,872 orthologous groups were identified across the six *T. atroviride* strains, of which 76% were shared by all strains. When considering subsets of closely related strains, 80% of orthologous groups were shared among strains I1237, P3041, P3080, and P3116, while 82% between strains MMS1295 and N1508. These results indicate that, although some differences in gene content exist among *T. atroviride* strains, the majority of genes are conserved. Orthologue groups were also identified in other studies spanning multiple *Trichoderma* species. For instance, a comparative analysis between *T. atroviride*, *T. virens*, and *T. reesei* revealed 15,682 orthologous groups, of which 7,915 were shared among the three species, representing 67% of the orthologues of *T. atroviride* (Kubicek et al., 2011). In a broader approach, 19,332 orthologous groups could be identified across 12 *Trichoderma* species, with 7,923 orthologs shared by at least two species within each clade, highlighting a much stronger interspecific divergence (Kubicek et al., 2019). Interestingly, the core genome of *Trichoderma* was found to be composed of 17% of genes absent in other fungal species (Steindorff et al., 2026).

While our comparative analysis indicates low overall genetic divergence between the studied strains, the main differences were observed in genes encoding NLR. A previous large-scale analysis across 195 fungal genomes showed that the genus *Trichoderma* has an expanded NLR repertoire compared to other fungi, particularly in mycoparasitic species such as *T. virens* (168 genes) and *T. atroviride* (135 genes), compared to the saprophytic *Trichoderma* species *T. reesei* (40 genes) (Dyrka et al., 2014). This expansion has been proposed to reflect specialized roles in environmental sensing and host recognition. In our dataset, HP strains displayed specific NLR orthologous groups absent from WP strains, including two putative fungal NLR uniquely detected in HP strains and upregulated in response to pathogen contact. Notably, the only two orthologous groups expanded in HP strains compared with WP strains also correspond to putative NLR-encoding genes. Together, these results suggest a potential role of NLR diversification in strain-specific responses to pathogens and may contribute to the contrasting transcriptomic profiles observed between genetically similar strains. Notably, WP strains retain a similar complement of genes associated with mycoparasitism but appear to exhibit a weaker transcriptional activation in response to pathogen contact, which may reflect differences in upstream perception or signal initiation rather than loss of effector capacity.

One of the mechanisms less expressed in WP strains is the production of cell wall–degrading enzymes, a key component of mycoparasitism that enables *Trichoderma* to degrade the prey cell wall, facilitating nutrient acquisition and host killing, as well as penetration and toxin delivery via *appressorium*-like structures. Chitinases (GH18 family) are the most abundant GH in *Trichoderma* (Gruber and Seidl-Seiboth, 2012), with up to 27 genes in *T. atroviride* and 33 in *T. virens* (Wang et al., 2021). Here, the six *T. atroviride* genomes contain between 28 and 31 GH18 genes. GH18 chitinases are commonly categorized (groups A, B, and C) based on their sequence features and domain architecture (Seidl et al., 2005).

Group A chitinases are ubiquitous across fungal genomes, with ascomycetes typically encoding an average of six (ranging from two in *Ustilago maydis* to 12 in *Fusarium oxysporum*). In our study, seven group A chitinases were identified in *T. atroviride*, consistent with previous findings (Wang et al., 2021). However, the presence of an additional group A chitinase in strains P3080 and P3116 is surprising, given the conserved nature of these enzymes. Since group A chitinases are highly conserved and not overrepresented in mycoparasitic fungi, they are hypothesized to play a limited role in mycoparasitism, instead contributing primarily to fungal development. Supporting this, the expression of chi18-5 (syn. ech42) has been linked to carbon starvation, suggesting a role in autolysis (Brunner et al., 2003). In our results, few group A chitinases were upregulated in response to pathogen contact and particularly in WP strains. While this expression occurs in response to physical contact and could be linked to parasitism, the known autolytic function of these chitinases, combined with the limited parasitic activity of these strains, suggests that it may instead reflect mycelial autolysis.

Group B chitinases exhibit considerable variability across fungal species. While most ascomycetes encode three or four, Hypocreales species average 10 (Wang et al., 2021). In *Trichoderma*, the mycoparasitic species *T. atroviride* and *T. virens* encode 12 group B chitinases, whereas the saprophyte *T. reesei* encodes only eight. This expansion of group B chitinases in *Trichoderma* is hypothesized to be linked to their mycoparasitic lifestyle, implying a more direct role in mycoparasitism (Seidl et al., 2005). In our study, 13 group B chitinases were always identified in *T. atroviride*. Many of these genes were significantly upregulated in response to contact with fungal prey, particularly in HP strains, supporting the hypothesis that these enzymes play a role in mycoparasitism.

Group C chitinases are highly variable, even within the *Trichoderma* genus, with *T. reesei*, *T. atroviride*, and *T. virens* encoding four, eight, and 13 group C chitinases, respectively (Wang et al., 2021). Our study also identified eight group C chitinases in *T. atroviride*, with three additional chitinases in strains MMS1295 and N1508. Group C chitinases share structural similarities with the α-subunit of the multimeric *Kluyveromyces lactis* killer toxin, which degrades the host cell wall to facilitate toxin penetration. Although the γ-subunit required for toxin synthesis has not been identified in *Trichoderma*, subgroup C chitinases may nonetheless contribute to host membrane permeabilization. Half of the eight enzymes in *T. atroviride* are upregulated in the presence of chitin, and all are upregulated in the presence of *B. cinerea* or its cell walls (Gruber et al., 2011). This result indicates that their expression is more linked to the interaction with the prey rather than solely for chitin consumption. However, in our study, subgroup C chitinases exhibited limited upregulation in response to contact with the tested pathogens, though certain subgroup C chitinases were significantly more expressed in HP strains. This differential expression suggests a potential, albeit nuanced, role for subgroup C chitinases in mycoparasitism, warranting further investigation.

Other fungal cell wall-degrading enzymes have also been linked to parasitism in *Trichoderma*. For instance, GH75 chitosanases are expanded in *Trichoderma* species compared to other ascomycetes, and further expanded in mycoparasitic *Trichoderma* relative to the saprophyte *T. reesei*. A similar pattern is observed for certain β-glucanases, such as those in the GH55 and GH64 families (Kubicek et al., 2011). In our study, these three hydrolase families were strongly upregulated in response to pathogen contact, particularly in HP strains. Our findings further reveal that other GH, including those involved in the degradation of cellulose, xylan, and other polysaccharides, are significantly upregulated in response to pathogen contact. Other fungal cell wall-degrading enzymes have also been linked to parasitism in *Trichoderma*. Several GH families (*e.g.*, GH27, GH30, GH54, GH55, GH64, GH65, GH71, GH75, GH79, GH92) are expanded in mycoparasitic *Trichoderma* species compared to the saprophyte *T. reesei* or other ascomycetes, some of which are known to participate in fungal cell wall degradation, while others are not (Gruber and Seidl-Seiboth, 2012; Kubicek et al., 2011). In our study, most of these GH genes were strongly upregulated in response to pathogen contact (Supplementary Fig. 13). Other CAZymes were also overexpressed in response to pathogen contact, including PL, CE, and AA families, suggesting a potential, yet understudied, role in mycoparasitism. Further investigation of these enzymes could provide valuable insights into their contribution to *Trichoderma* parasitic interactions.

Peptidases and proteases may also play a critical role in degrading fungal cell wall integrity. Mycoparasitic *Trichoderma* species exhibit a markedly expanded repertoire of peptidase-encoding genes: *T. reesei* contains only two genes with protease-associated PFAM domains, whereas *T. virens* and *T. atroviride* possess 28 and 23, respectively (Kubicek et al., 2011). Transcriptomic analyses have consistently demonstrated the upsurge in peptidase expression in response to contact between *T. atroviride* and its prey (Chen et al., 2023; Seidl et al., 2009b). In our study, several peptidases were also upregulated in response to pathogen contact, particularly those belonging to the S08 family. The S08 peptidases represent one of the most expanded paralog groups in *Trichoderma*: *T. reesei* encodes 10 (comparable to other fungi), while *T. virens* and *T. atroviride* encode 22 and 36, respectively, based on PFAM domain analysis (Kubicek et al., 2011). Notably, the expression of secreted peptidase is more important in HP compared to WP strains, particularly among serine peptidases. The low expression of these cell wall–degrading enzymes essential for mycoparasitism in *Trichoderma* may therefore explain the reduced antagonistic activity observed in WP strains.

Another important mechanism of *Trichoderma* mycoparasitism is the production of various specialized metabolites, some of which display antimicrobial activity (Hermosa et al., 2014). A diversity of 39 PKS have been identified in various *Trichoderma* species which contain between 9 and 20, including 18 in *T. atroviride* (Kubicek et al., 2019, 2011). In our study, 17 or 18 PKS were identified across the six *T. atroviride* strains. The number of NPRS coding genes also varies between and within *Trichoderma* species (11-29 genes, including 16 in *T. atroviride*). In our study, 13 NRPS and 10 NRPS-like were identified. Additionally, while a range of 6 to 12 terpene synthases have been identified in *Trichoderma* species, the strains of our study count for 12 to 14. These results show that the number of BGC varies both among *Trichoderma* species and within a single species. A substantial proportion of biosynthetic genes (mostly PKS, NRPS and isocyanide biosynthetic genes) are upregulated in response to pathogen contact. Moreover, 72% of these genes are differentially expressed between WP and HP strains. These results indicate that HP strains do not exhibit a global increase in the expression of biosynthetic genes, but rather distinct patterns across individual genes that could be translated into different metabolic profiles between WP and HP strains, with specific compounds upregulated and/or *de novo* induced in *T. atroviride* when confronted to the pathogens. As such metabolic regulation is rather complexe and specific (Arora et al., 2020), further investigations of strains metabolomes would help identifying candidate molecules with potential biofungicide properties.

In addition to attack-related mechanisms in *Trichoderma*, our study also highlights the involvement of defense mechanisms in mycoparasitic performance. Among the genes more highly expressed in *T. atroviride* HP strains, several are involved in detoxification and stress-response pathways, including multiple components of the HOG signaling pathway. They include key elements of the Sln1 branch and downstream transcription factors regulating the expression of stress-responsive genes. Functional studies in other *Trichoderma* species support the importance of this pathway in stress adaptation. This is illustrated by the roles of Ypd1 in *T. reesei* and Hog1 in *T. harzianum* in responses to osmotic, oxidative, and thermal stresses, as well as in maintaining cell wall integrity. Furthermore, in *T. harzianum* HOG1 is essential for antagonism against *Phoma betae* and *Colletotrichum acutatum* (Delgado-Jarana et al., 2006; Wang et al., 2018). The combined enhanced activation of the Sln1–Hog1 pathway in HP strains may contribute to their improved stress tolerance and, consequently, to their increased antagonistic capacity. In HP strains, the higher expression of genes of the HOG pathway is accompanied by a similar pattern of ROS detoxification-related genes (*e.g.*, superoxide dismutase, catalase, peroxidase, peroxiredoxin). In addition, these enzymes, along with others involved in xenobiotic detoxification, are induced in response to pathogen contact, consistent with the reported induction of a glutathione peroxidase in *T. atroviride* during interaction with *Armillaria ostoyae* (Chen et al., 2023).

Consistent with the enhanced expression of detoxification pathways, the peroxisome, an intracellular organelle involved in oxidative stress management, also appears to be differentially regulated in HP strains. The peroxisome hosts several ROS-detoxifying enzymes, including superoxide dismutases and catalases. Its biogenesis is mediated by a set of highly conserved proteins known as peroxins (PEX) (Maruyama and Kitamoto, 2013). Genes belonging to the PEX family have been shown to be upregulated in *T. atroviride* in response to contact with *A. ostoya*e (Chen et al., 2023), as well as in *T. asperellum* during interaction with the pinewood nematode *Bursaphelenchus xylophilus* (Chen et al., 2024). In our study, PEX11, a gene involved in peroxisome proliferation (Escaño et al., 2009), was upregulated in response to pathogen contact in all *T. atroviride* strains, suggesting a potential increase in peroxisome abundance. Notably, six other PEX genes are specifically overexpressed in response to pathogen contact in HP strains only. These results suggest an increased peroxisomal activity and proliferation in HP strains during pathogen interaction. Beyond their role in ROS detoxification, peroxisomes are also involved in fatty acid catabolism through β-oxidation. Several genes associated with this pathway were more highly expressed in HP strains. Upregulation of β-oxidation-related genes in response to pathogen contact has previously been reported in *T. atroviride* against *B. cinerea* and *R. solani* (Seidl et al., 2009b). It has been proposed that this metabolic shift may be linked to appressorium formation, as observed in *Pyricularia oryzae* (Oh et al., 2008). In this species, β-oxidation contributes to glycerol accumulation, which generates the turgor pressure required for host penetration, although such a mechanism has not yet been demonstrated in *Trichoderma*. Consistent with this, peroxisomes have been shown to be essential for appressorium-mediated host penetration in *Colletotrichum lagenarium* (Kimura et al., 2001). Another key function of peroxisomes in filamentous fungi is the formation of Woronin bodies, organelles that ensure hyphal compartmentalization and prevent cytoplasmic leakage following mechanical damage. Woronin bodies have been associated with multiple biological processes, including hyphal growth, stress tolerance, and pathogenicity (Maruyama and Kitamoto, 2013). HEX1, the major structural component of Woronin bodies, is upregulated in *T. atroviride* during growth on fungal cell walls (Marra et al., 2006). In *T. simmonsii*, HEX1 has been implicated in preventing cytoplasmic leakage, in fungal growth and mycoparasitism performance against fungi and oomycetes (Pedrero-Méndez et al., 2025). Consistently, in our study, HEX1 is more highly expressed in HP strains compared to WP strains. Overall, these coordinated responses point to a reinforced detoxification capacity in HP strains, integrating HOG signaling, ROS-scavenging systems, and peroxisome-associated processes, which likely contributes to improved stress tolerance during mycoparasitism.

Fungal interaction with other organisms is often mediated by a diverse repertoire of small secreted proteins enriched in cysteine residues, which are thought to act as key effectors in interspecies communication. A comparative genomic analysis across 12 *Trichoderma* species identified between 39 and 125 small secreted cysteine-rich proteins (SSCPs) (Kubicek et al., 2019). These results highlight substantial variability in SSCP content both across *Trichoderma* species and also within *T. atroviride* strains in our study (between 113 and 121). Another study identified 86 effector-like proteins in *T. atroviride*, as well as 84 in *T. virens* and only 63 in the saprophytic *T. reesei* (Guzmán-Guzmán et al., 2017). The authors showed that several of these genes were upregulated in response to contact with *R. solani* as well as during interaction with *Arabidopsis thaliana* roots. In our analysis, among the 113 SSCPs identified, 23 were consistently upregulated in response to pathogen contact across all strains, while 17 were specifically upregulated by HP strains and 22 by WP strains. The integration of genomic and transcriptomic data thus provides a powerful approach to identify SSCP repertoires and to prioritize candidates based on their expression profiles during interaction. However, at this stage, the precise biological functions of these SSCP cannot be inferred and will require targeted functional validation. Several effectors have already been functionally validated in *Trichoderma*, including Tal6, a LysM effector involved in mycoparasitism and plant association (Romero-Contreras et al., 2019) or a hydrophobin in *T. virens* which enhance mycoparasitic performance against *R. solani* as well as promote root colonization of *A. thaliana* (Guzmán-Guzmán et al., 2017). These results suggest that these effector-like proteins may be involved in *Trichoderma*–prey interactions, and that the differences observed between WP and HP strains could contribute to the variation in antagonistic activity between these strains.

## 5 Conclusion

This analysis provides initial insights into the molecular basis of mycoparasitic performance. Based on our results, the performance of the four HP strains can be attributed, at least in part, to three distinct expression patterns (Fig. 11): (i) mechanisms linked to mycoparasitism, induced in all strains in response to pathogen contact but more strongly upregulated in HP strains (*e.g.*, cell wall-degrading enzymes); (ii) HP-specific mechanisms, overexpressed in HP strains compared to WP strains but not induced by pathogen contact (*e.g.*, ROS detoxification and defense pathways); and (iii) strain-specific mechanisms involving divergent gene sets between WP and HP strains (*e.g.*, specialized metabolism and effector-like proteins). All genes involved in these mechanisms are present in the genomes of WP strains but are expressed at lower levels compared to HP strains. This difference may therefore result from a reduced ability of WP strains to perceive their prey. This hypothesis is consistent with the loss of certain NLR receptor genes in the genomes of WP strains.

**Fig. 11.**
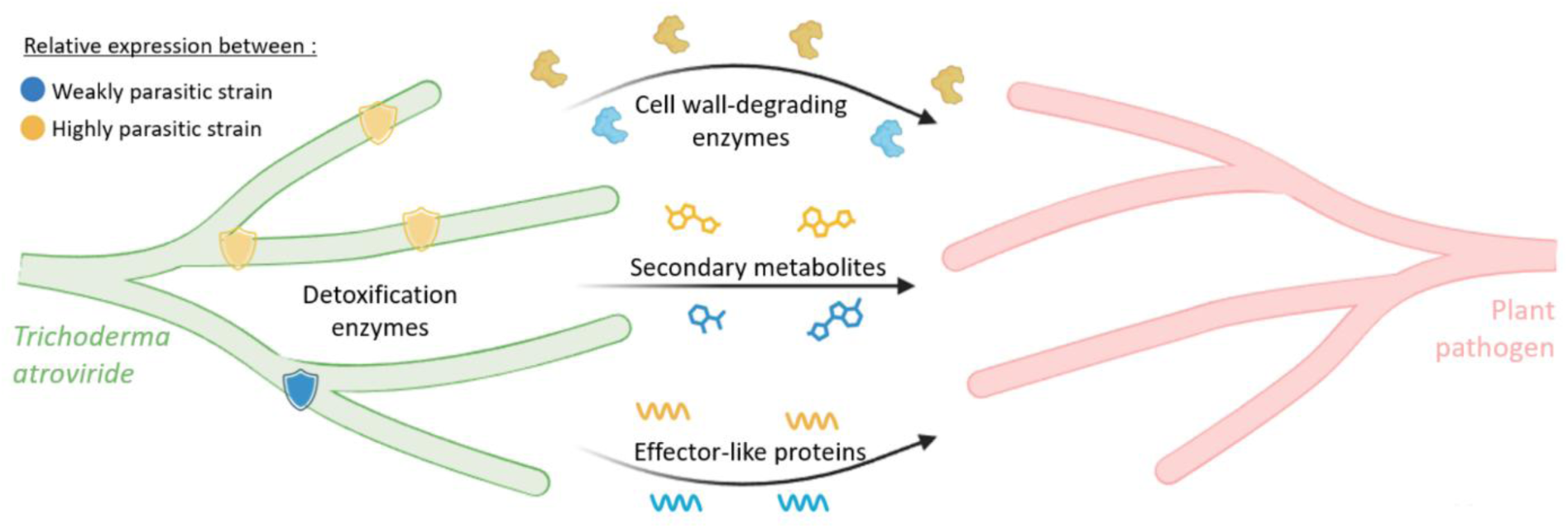
Schematic representation of differences in gene expression between WP and HP strains. Each item in the schematic represents genes that are overexpressed in WP strains (blue) or in HP strains (yellow) after contact with a pathogenic agent.

Together, these findings underscore the necessity of extensive functional validation. For detoxification pathway, biosynthesis of specialized metabolites and NLR receptors, targeted mutagenesis of key actors will be essential to evaluate their specific contributions. Conversely, for effector-like proteins and cell wall-degrading enzymes, functional redundancy often masks phenotypes in single-gene knockouts. Overexpression strategies or targeting master regulators may be more effective. The identification of these master regulators could be facilitated by leveraging WGCNA to identify hub genes within expression modules that are highly correlated with both mycoparasitic traits and module membership (Langfelder and Horvath, 2008). Specifically, transcription factors acting as hubs in modules enriched for polysaccharide-degrading enzymes (*e.g.*, module 2) could represent key regulators of these enzymatic families. Furthermore, the integration of Gene Regulatory Network (GRN) modeling, which combines transcriptomic data with promoter motif analysis, could further streamline the identification of master regulators, as previously demonstrated in *T. atroviride* (Olivares-Yañez et al., 2021).

Comparative genomics revealed only limited differences between *T. atroviride* strains but the increasing availability of high-quality genomes offers new opportunities to explore the genetic basis of phenotypic variation in *Trichoderma* (Steindorff et al., 2026). While ortholog-based comparisons revealed relatively few differences among *T. atroviride* strains, genome-wide association studies (GWAS) represent a promising complementary approach. By analyzing a larger and more diverse collection of isolates, it should become possible to associate SNP (Single Nucleotide Polymorphism) with quantitative traits such as mycoparasitic performance. Ultimately, combining functional validation, regulatory analyses, and expanded genomic resources will be key to identifying robust molecular markers associated with mycoparasitic performance. This could allow rapid selection of efficient strains and, if sexual reproduction can be harnessed as in *T. reesei* (Seidl et al., 2009a), support marker-assisted breeding strategies to accelerate the development of improved biocontrol agents.

## CRediT Authorship Contribution Statement

**Etienne Brémand**: Conceptualization, Investigation, Methodology, Software, Formal analysis, Visualization, Writing – original draft. **Franck Bastide**: Conceptualization, Investigation, Methodology, Writing – review & editing. **Justine Colou**: Conceptualization, Investigation, Methodology, Writing – review & editing. **Nicolas Denancé**: Conceptualization, Writing – review & editing. **Séverine Boisard**: Conceptualization, Writing – review & editing. **Nicolas Ruiz**: Conceptualization, Writing – review & editing. **Samuel Bertrand**: Conceptualization, Writing – review & editing. **Muriel Marchi**: Methodology, Writing – review & editing. **Jérôme Verdier**: Software, Writing – review & editing. **Thomas Guillemette**: Conceptualization, Supervision, Project administration, Funding acquisition, Writing – review & editing.

## Funding Sources

This work was supported by the University of Angers and Angers Loire Métropole, which co-funded the PhD fellowship associated with this publication.

## Acknowledgments

We are grateful to the ANAN platform (SFR QUASAV), especially Muriel Bahut, and to the PACEM platform (SFR ICAT), especially Jérôme Cayon, for technical and material support. We also acknowledge the GenOuest bioinformatics core facility (https://www.genouest.org) for providing computational infrastructure.

## Declaration Of Competing Interests

The authors declare no conflict of interest.

## Appendix A. Supplementary Material

Supplementary Figures and Supplementary Tables

Supplementary Dataset 1. Functional annotation and transcriptional analysis of each gene in the *Trichoderma atroviride* N1508 genome. For each gene, the table details its presence in a WGCNA module, the results of the differential expression analysis, and the results of all functional annotations (EggNOG, CAZyme, BGC, detoxification genes, effectors).

Supplementary Dataset 2. Orthologue groups among the six *Trichoderma atroviride* strains. This file details all orthologue groups identified among the six *T. atroviride* strains (including those not shared between strains). For each group, the gene names are detailed for all six strains.

## Data Availability

All data are accessible at the National Center for Biotechnology Information (NCBI) under BioProject accession PRJEB108439 (ncbi.nlm.nih.gov/bioproject/PRJEB108439). This project includes raw DNA and RNA sequencing data, as well as genome assemblies and annotations for the *T. atroviride* strains. A comprehensive list of all gene identifiers and annotations discussed in this study is provided in Supplementary Dataset 1, and the results of the orthologue analysis are provided in Supplementary Dataset 2. All scripts used for the genomic and transcriptomic analyses are available on GitHub (github.com/ebremand/Trichoderma-Mycoparasitic-Performance/tree/main) and have been deposited on Zenodo (doi.org/10.5281/zenodo.19694549).

## Notes

https://www.ncbi.nlm.nih.gov/bioproject/PRJEB108439

https://github.com/ebremand/Trichoderma-Mycoparasitic-Performance/tree/main

### Competing Interest Statement

The authors have declared no competing interest.

