## Supplementary Figure and Supplementary Table for "Molecular Basis of Mycoparasitic Performance: Genomic and Transcriptomic Comparison of Contrasting *Trichoderma atroviride* Strains"

Supplementary Table 1. Genomes used for the genomic distance tree (Figure 2).
At the top, the six *Trichoderma atroviride* strains from this study. In the middle, the ten other *T. atroviride* strains. At the bottom, one *Trichoderma gamsii* strain used to root the tree.

| GenBank | Species | Strain | Size (Mb) | Scaffold | Contigs | Level | Release date | Contig N50 (kb) | Scaffold N50 (kb) |
|---|---|---|---|---|---|---|---|---|---|
| GCA_980757495.1 | *Trichoderma atroviride* | I1237 | 37.20 | 11 | 11 | Contig | apr-2026 | 5,599 | 5,599 |
| GCA_980757575.1 | *Trichoderma atroviride* | P3041 | 37.24 | 9 | 9 | Contig | apr-2026 | 5,623 | 5,623 |
| GCA_980757555.1 | *Trichoderma atroviride* | P3080 | 37.52 | 52 | 52 | Contig | apr-2026 | 1,961 | 1,961 |
| GCA_980753795.1 | *Trichoderma atroviride* | P3116 | 37.45 | 33 | 33 | Contig | apr-2026 | 3,744 | 3,744 |
| GCA_982374855.1 | *Trichoderma atroviride* | MMS1295 | 36.74 | 22 | 22 | Contig | apr-2026 | 3,084 | 3,084 |
| GCA_982374845.1 | *Trichoderma atroviride* | N1508 | 36.93 | 16 | 16 | Contig | apr-2026 | 3,39 | 3,39 |
| GCA_051557965.1 | *Trichoderma atroviride* | B7 | 37.96 | 52 | 52 | Contig | jul-25 | 3,455 | 3,455 |
| GCA_041053265.1 | *Trichoderma atroviride* | CGMCC 40488 | 36.34 | 8 | 20 | Chromosome | aug-24 | 2,647 | 5,43 |
| GCA_040333345.1 | *Trichoderma atroviride* | F020 | 36.45 | 7 | 15 | Scaffold | jun-24 | 3,047 | 5,495 |
| GCA_028554805.1 | *Trichoderma atroviride* | SC1 | 35.76 | 603 | 603 | Contig | feb-23 | 312 | 312 |
| GCA_020647795.1 | *Trichoderma atroviride* | P1 | 37.30 | 7 | 7 | Complete | oct-21 | 5,658 | 5,658 |
| GCA_020466355.1 | *Trichoderma atroviride* | CG 6828 | 36.66 | 37 | 37 | Contig | oct-21 | 1,578 | 1,578 |
| GCA_019297715.1 | *Trichoderma atroviride* | IMI 206040 | 36.16 | 12 | 12 | Contig | jul-21 | 5,621 | 5,621 |
| GCA_002916895.1 | *Trichoderma atroviride* | LY357 | 35.90 | 637 | 637 | Contig | feb-18 | 100 | 100 |
| GCA_001599035.1 | *Trichoderma atroviride* | JCM 9410 | 37.32 | 23 | 240 | Scaffold | mar-16 | 532 | 5,619 |
| GCA_000963795.1 | *Trichoderma atroviride* | XS2015 | 36.40 | 131 | 357 | Scaffold | mar-15 | 279 | 2,105 |
| GCA_978148955.1 | *Trichoderma gamsii* | T035 | 38.8 | 16 | 28 | Scaffold | mar-26 | 3,095 | 7,178 |

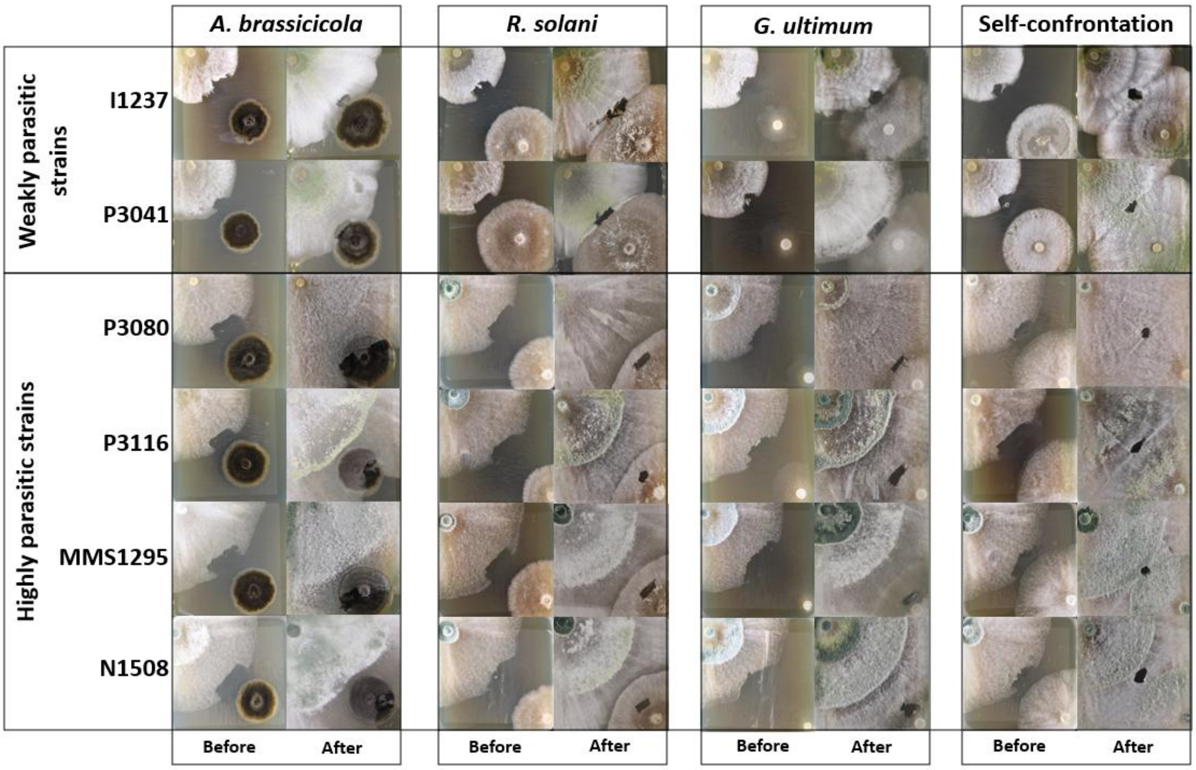


Supplementary Fig 1. Photographs of the *in vitro* confrontation assays used for RNA-seq analysis. The six *Trichoderma atroviride* strains are shown in rows, and the four confrontation conditions are shown in columns, with photographs representing the stages before and after contact between the two fungi. In each image, *T. atroviride* is located in the upper right and the pathogen in the lower left. The holes in the *T. atroviride* mycelium correspond to the sampling sites used for RNA-seq.

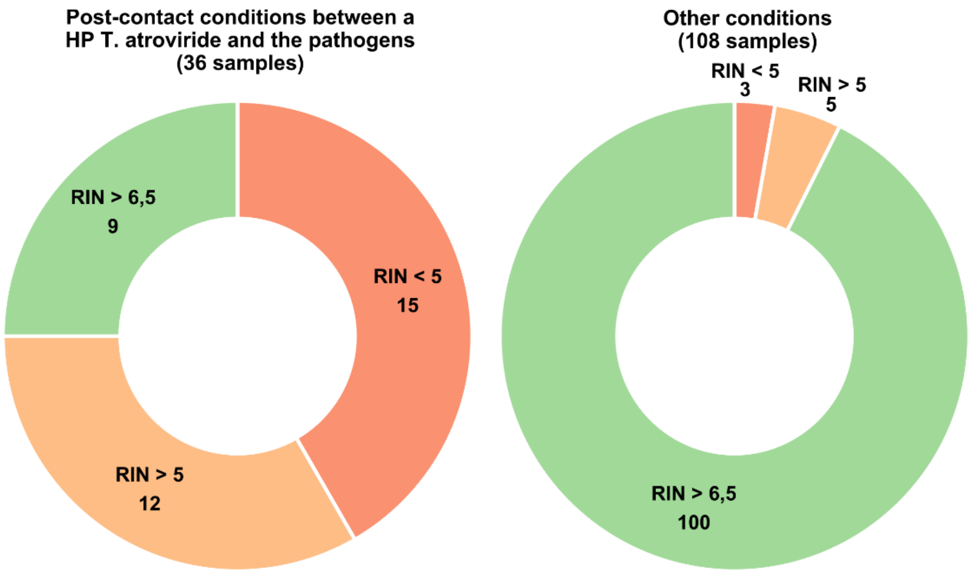


Supplementary Fig. 2. RNA Integrity Number (RIN) of all samples from the first RNA extraction. Left, samples after contact between a HP *Trichoderma atroviride* strain and the pathogens. Right, all pre-contact conditions, post-contact conditions for WP *T. atroviride* strains, and post-contact conditions between *T. atroviride* strains themselves.

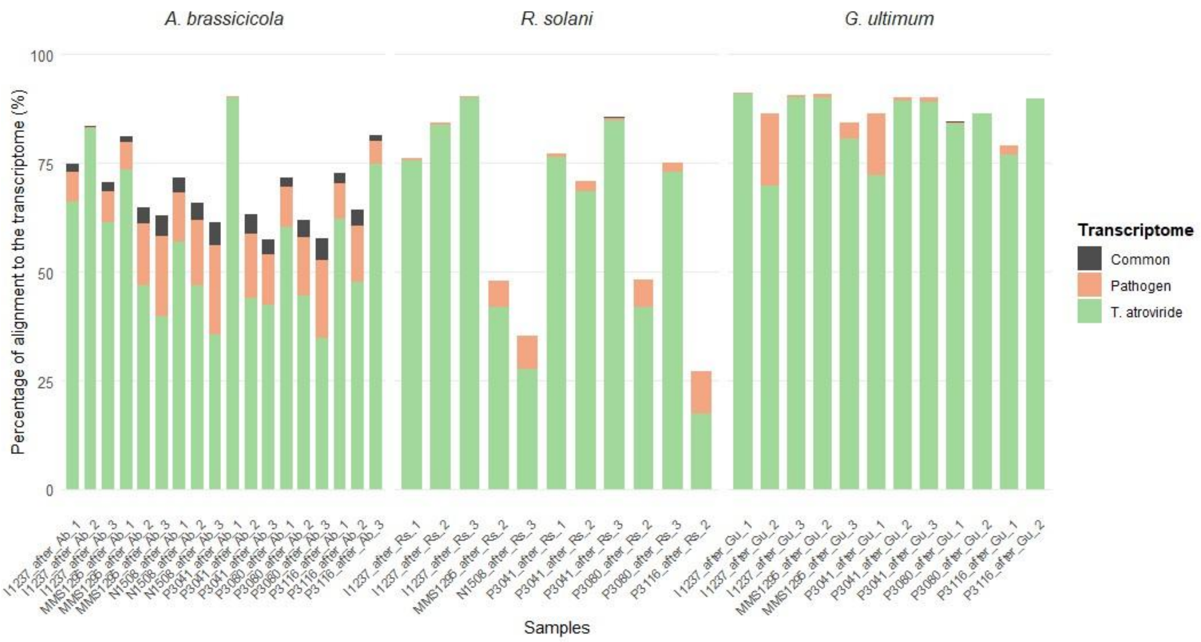


Supplementary Fig. 3. Percentage of read alignment to reference transcriptomes.
Percentage of alignment of samples after pathogen contact to the transcriptomes of *T. atroviride* N1508, the pathogen (*Alternaria brassicicola*, *Rhizoctonia solani*, *Globisporangium ultimum*), or both.

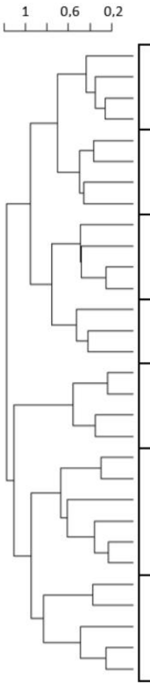


| | Weakly parasitic *T. atroviride* strains | | | | | | | | Highly parasitic *T. atroviride* strains | | | | | | | | | | | | | | | | Number of genes |
|---|---|---|---|---|---|---|---|---|---|---|---|---|---|---|---|---|---|---|---|---|---|---|---|---|---|
| | I1237 / P3041 | | | | | | | | P3080 / P3116 | | | | | | | | MMS1295 / N1508 | | | | | | | | |
| | Before contact | | | | After contact | | | | Before contact | | | | After contact | | | | Before contact | | | | After contact | | | | |
| | Ab | Rs | Gu | Ta | Ab | Rs | Gu | Ta | Ab | Rs | Gu | Ta | Ab | Rs | Gu | Ta | Ab | Rs | Gu | Ta | Ab | Rs | Gu | Ta | |
| EM 1.1 | -0,05 | -0,05 | -0,05 | 0,00 | 0,02 | 0,04 | -0,01 | 0,10 | -0,06 | -0,07 | -0,03 | 0,02 | -0,01 | 0,08 | 0,01 | 0,16 | -0,07 | -0,08 | -0,05 | -0,02 | 0,00 | 0,04 | 0,06 | 0,14 | 365 |
| EM 1.2 | -0,05 | -0,05 | -0,09 | -0,02 | 0,04 | 0,08 | -0,05 | 0,07 | -0,02 | -0,05 | -0,06 | -0,01 | -0,05 | -0,10 | 0,09 | 0,11 | -0,04 | -0,06 | 0,03 | 0,08 | -0,04 | -0,07 | 0,11 | 0,13 | 140 |
| EM 1.3 | -0,03 | -0,03 | -0,04 | 0,02 | 0,10 | 0,14 | -0,01 | 0,15 | -0,04 | -0,06 | -0,06 | -0,04 | 0,00 | -0,04 | -0,01 | 0,06 | -0,06 | -0,06 | -0,05 | 0,02 | -0,01 | -0,03 | 0,00 | 0,04 | 638 |
| EM 1.4 | -0,09 | -0,10 | -0,11 | -0,02 | 0,14 | 0,12 | -0,08 | 0,10 | -0,04 | -0,05 | -0,03 | 0,01 | 0,04 | 0,00 | 0,00 | 0,04 | -0,04 | -0,04 | 0,02 | 0,03 | 0,02 | 0,05 | 0,01 | 0,06 | 147 |
| EM 2.1 | -0,07 | -0,07 | -0,08 | -0,05 | 0,10 | 0,10 | -0,06 | 0,01 | -0,04 | -0,05 | -0,06 | -0,04 | 0,11 | 0,00 | -0,02 | -0,01 | -0,01 | -0,02 | -0,01 | 0,02 | 0,13 | 0,11 | 0,02 | 0,08 | 81 |
| EM 2.2 | -0,02 | -0,04 | -0,06 | -0,05 | 0,07 | 0,03 | -0,04 | -0,05 | 0,00 | -0,05 | -0,06 | -0,04 | 0,25 | 0,11 | -0,01 | -0,04 | -0,02 | -0,05 | -0,05 | -0,04 | 0,22 | 0,10 | -0,01 | -0,04 | 211 |
| EM 2.3 | -0,04 | -0,05 | -0,05 | -0,04 | 0,02 | 0,16 | -0,04 | -0,03 | -0,01 | -0,01 | -0,01 | -0,01 | 0,08 | 0,27 | 0,00 | -0,03 | -0,04 | -0,04 | -0,04 | -0,03 | 0,03 | 0,39 | -0,03 | -0,05 | 80 |
| EM 2.4 | -0,10 | -0,12 | -0,12 | -0,04 | 0,08 | 0,05 | -0,12 | 0,00 | -0,05 | -0,04 | 0,01 | 0,03 | 0,05 | 0,12 | -0,02 | -0,02 | 0,01 | 0,00 | 0,08 | 0,09 | 0,04 | 0,19 | 0,06 | 0,04 | 79 |
| EM 3.1 | -0,07 | -0,07 | -0,08 | -0,07 | -0,06 | -0,05 | -0,07 | -0,06 | -0,05 | -0,06 | -0,06 | -0,05 | -0,02 | -0,05 | -0,03 | -0,04 | 0,11 | 0,11 | 0,12 | 0,13 | 0,16 | 0,12 | 0,15 | 0,14 | 639 |
| EM 3.2 | -0,07 | -0,08 | -0,10 | -0,09 | -0,09 | -0,10 | -0,12 | -0,13 | 0,06 | 0,05 | 0,06 | 0,05 | 0,10 | 0,14 | 0,03 | -0,04 | 0,06 | 0,06 | 0,07 | 0,04 | 0,10 | 0,15 | 0,06 | 0,00 | 276 |
| EM 3.3 | -0,12 | -0,12 | -0,14 | -0,11 | -0,08 | -0,07 | -0,10 | -0,08 | 0,08 | 0,05 | 0,02 | 0,06 | 0,07 | 0,05 | 0,14 | 0,08 | 0,04 | 0,02 | 0,04 | 0,05 | 0,06 | 0,03 | 0,11 | 0,10 | 470 |
| EM 3.4 | -0,11 | -0,11 | -0,16 | -0,07 | 0,03 | 0,01 | -0,11 | -0,02 | 0,03 | 0,01 | -0,01 | 0,04 | -0,03 | -0,08 | 0,11 | 0,07 | 0,07 | 0,02 | 0,08 | 0,10 | -0,04 | -0,07 | 0,12 | 0,12 | 208 |
| EM 4.1 | -0,02 | 0,01 | -0,04 | 0,00 | -0,07 | -0,01 | 0,06 | -0,05 | 0,05 | 0,06 | 0,04 | 0,05 | -0,16 | -0,13 | 0,09 | 0,02 | 0,07 | 0,04 | 0,05 | 0,05 | -0,16 | -0,21 | 0,05 | 0,06 | 83 |
| EM 4.2 | -0,08 | -0,07 | -0,10 | -0,05 | 0,07 | 0,00 | -0,02 | -0,10 | 0,01 | 0,02 | 0,01 | 0,02 | -0,02 | -0,11 | 0,03 | -0,04 | 0,10 | 0,06 | 0,12 | 0,11 | -0,04 | -0,15 | 0,07 | 0,04 | 112 |
| EM 4.3 | -0,04 | -0,07 | -0,08 | 0,05 | -0,01 | 0,00 | -0,07 | 0,00 | -0,02 | 0,01 | 0,09 | 0,10 | -0,09 | 0,02 | -0,03 | -0,03 | 0,03 | 0,02 | 0,13 | 0,11 | -0,11 | -0,05 | 0,02 | 0,02 | 42 |
| EM 5.1 | 0,03 | 0,00 | 0,02 | 0,01 | -0,07 | -0,08 | -0,05 | -0,08 | 0,04 | 0,06 | 0,08 | 0,02 | 0,04 | 0,09 | -0,04 | -0,15 | 0,04 | 0,06 | 0,04 | 0,02 | 0,06 | 0,13 | -0,01 | -0,14 | 66 |
| EM 5.2 | 0,11 | 0,08 | 0,10 | 0,03 | -0,07 | -0,08 | -0,05 | -0,07 | 0,01 | 0,02 | 0,02 | -0,01 | 0,00 | 0,07 | -0,05 | -0,12 | 0,04 | 0,04 | 0,01 | -0,03 | 0,03 | 0,11 | -0,02 | -0,09 | 339 |
| EM 5.3 | -0,02 | 0,00 | 0,02 | -0,01 | -0,07 | -0,09 | -0,03 | -0,10 | 0,03 | 0,09 | 0,10 | 0,03 | -0,03 | -0,01 | -0,05 | -0,12 | 0,09 | 0,12 | 0,10 | 0,06 | -0,02 | -0,02 | -0,06 | -0,10 | 198 |
| EM 5.4 | -0,01 | 0,03 | 0,07 | 0,01 | -0,05 | -0,05 | 0,09 | -0,03 | 0,01 | 0,07 | 0,06 | -0,02 | -0,03 | -0,07 | -0,14 | -0,07 | 0,07 | 0,12 | 0,04 | 0,00 | -0,02 | -0,05 | -0,15 | -0,10 | 259 |
| EM 6.1 | 0,07 | 0,08 | 0,07 | 0,08 | 0,13 | 0,14 | 0,08 | 0,14 | -0,06 | -0,08 | -0,09 | -0,06 | 0,00 | -0,03 | -0,04 | 0,02 | -0,10 | -0,10 | -0,10 | -0,08 | -0,03 | -0,06 | -0,04 | -0,04 | 628 |
| EM 6.2 | 0,16 | 0,15 | 0,14 | 0,11 | 0,03 | 0,04 | 0,06 | 0,08 | -0,06 | -0,06 | -0,06 | -0,06 | -0,04 | -0,01 | -0,06 | -0,06 | -0,06 | -0,06 | -0,07 | -0,07 | -0,02 | -0,02 | -0,04 | -0,07 | 197 |
| EM 6.3 | -0,03 | 0,01 | 0,03 | -0,01 | 0,15 | 0,11 | 0,13 | 0,01 | -0,03 | -0,06 | -0,07 | -0,06 | 0,11 | -0,04 | -0,02 | -0,02 | -0,05 | -0,06 | -0,06 | -0,04 | 0,05 | -0,07 | -0,04 | -0,04 | 290 |
| EM 6.4 | 0,11 | 0,14 | 0,13 | 0,04 | -0,08 | -0,03 | 0,11 | -0,03 | 0,00 | 0,03 | 0,02 | -0,04 | -0,11 | -0,06 | -0,02 | -0,09 | 0,04 | 0,05 | 0,00 | -0,04 | -0,10 | -0,09 | -0,04 | -0,07 | 90 |
| EM 6.5 | 0,09 | 0,14 | 0,14 | 0,03 | 0,07 | 0,06 | 0,15 | 0,02 | -0,03 | -0,03 | -0,04 | -0,07 | 0,00 | -0,04 | -0,04 | -0,06 | -0,04 | -0,04 | -0,07 | -0,07 | -0,04 | -0,10 | -0,06 | -0,10 | 71 |
| EM 6.6 | 0,01 | 0,07 | 0,09 | -0,01 | 0,05 | 0,01 | 0,14 | -0,05 | -0,01 | 0,01 | 0,01 | -0,04 | 0,01 | -0,06 | -0,07 | -0,08 | 0,03 | 0,04 | 0,00 | -0,03 | -0,03 | -0,11 | -0,09 | -0,08 | 292 |
| EM 7.1 | -0,06 | -0,06 | -0,07 | -0,06 | 0,00 | -0,01 | -0,04 | -0,04 | 0,13 | 0,10 | 0,08 | 0,11 | 0,15 | 0,11 | 0,16 | 0,09 | -0,09 | -0,10 | -0,09 | -0,08 | -0,02 | -0,08 | -0,04 | -0,06 | 136 |
| EM 7.2 | 0,03 | 0,04 | 0,02 | 0,02 | 0,09 | 0,09 | 0,06 | 0,06 | 0,06 | 0,03 | 0,00 | 0,03 | 0,10 | 0,02 | 0,09 | 0,08 | -0,14 | -0,15 | -0,14 | -0,12 | -0,07 | -0,14 | -0,08 | -0,10 | 173 |
| EM 7.3 | 0,10 | 0,08 | 0,05 | 0,08 | -0,03 | 0,00 | -0,03 | 0,07 | -0,01 | -0,05 | -0,04 | 0,00 | -0,10 | 0,01 | 0,00 | 0,06 | -0,04 | -0,06 | -0,05 | -0,05 | -0,08 | -0,04 | 0,06 | 0,08 | 157 |
| EM 7.4 | 0,03 | 0,02 | 0,04 | 0,02 | -0,11 | -0,06 | 0,07 | 0,07 | 0,06 | 0,03 | 0,06 | 0,09 | -0,11 | 0,00 | 0,04 | 0,05 | -0,06 | -0,01 | -0,07 | -0,05 | -0,09 | -0,06 | -0,01 | 0,02 | 84 |
| EM 7.5 | 0,04 | 0,03 | 0,03 | 0,03 | -0,06 | 0,01 | 0,05 | 0,15 | 0,01 | -0,03 | -0,01 | 0,04 | -0,08 | 0,04 | 0,03 | 0,12 | -0,10 | -0,08 | -0,11 | -0,08 | -0,07 | -0,03 | 0,00 | 0,08 | 138 |

Supplementary Fig. 4. Heatmap of module eigengene expression according to *Trichoderma atroviride* strain, timepoint (before or after contact), and the pathogen used in confrontation (Ab = *Alternaria brassicicola*, Rs = *Rhizoctonia solani*, Gu = *Globisporangium ultimum*, Ta = confrontation of the *T. atroviride* strain with itself). On the left, dendrogram of WGCNA modules based on gene expression similarity, with the scale representing module distance (1 – correlation). On the right, the number of genes in each expression module (EM).

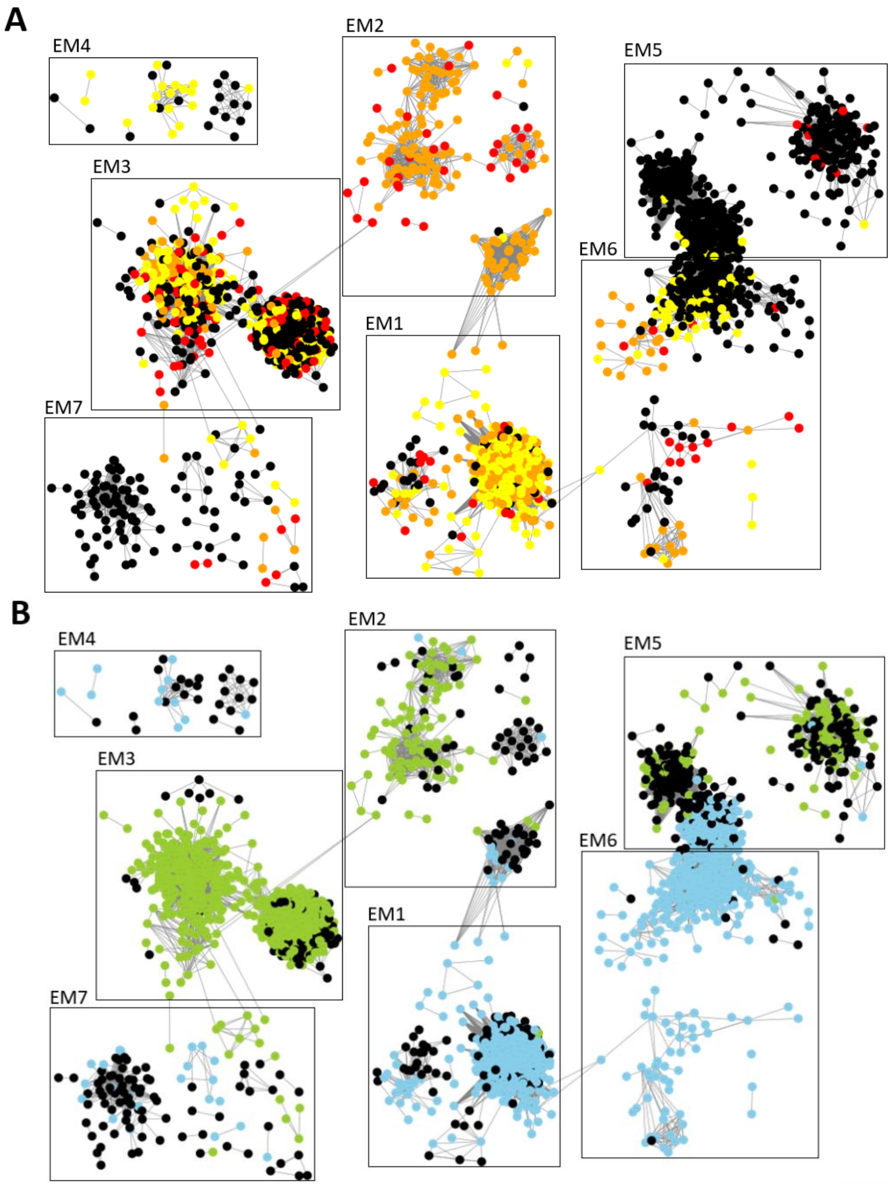


Supplementary Fig. 5: Relationship between the gene co-expression network and differential expression analysis.

The network was visualized using Cytoscape, considering only correlations above 0.15. Rectangles represent the seven expression module groups (EM). At the top (A), genes are colored according to their expression changes in response to pathogen contact: yellow indicates genes upregulated in response to contact in WP strains, red indicates genes upregulated in HP strains, and orange indicates genes upregulated in both strain types. At the bottom (B), genes are colored according to their differential expression after pathogen contact between HP and WP strains: green indicates genes more highly expressed in HP strains, and blue indicates genes more highly expressed in WP strains. In black, non-differentially expressed genes.

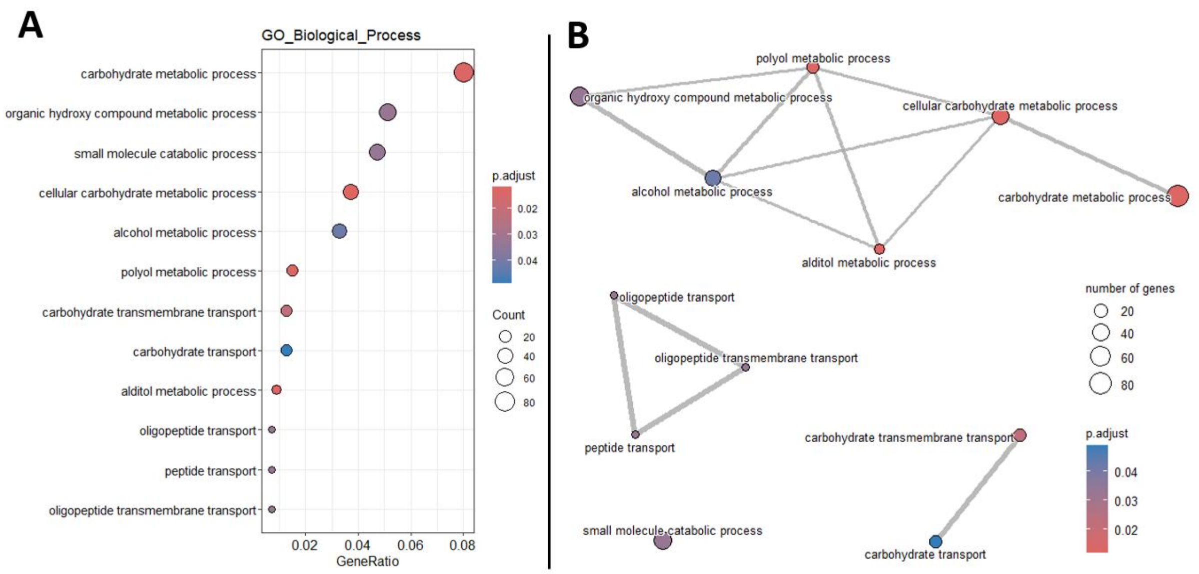


Supplementary Fig. 6. Functional enrichment of genes from expression module 1. Expression modules 1.1, 1.2, 1.3, and 1.4 comprise 1,290 genes that are more highly expressed in the six *T. atroviride* strains following self-confrontation and, in the weakly parasitic strains, after confrontation with *A. brassicicola* and *R. solani*. Dot plot (A) and Enrichment Map (B) of the 30 most significantly enriched GO (Gene Ontology) biological processes.

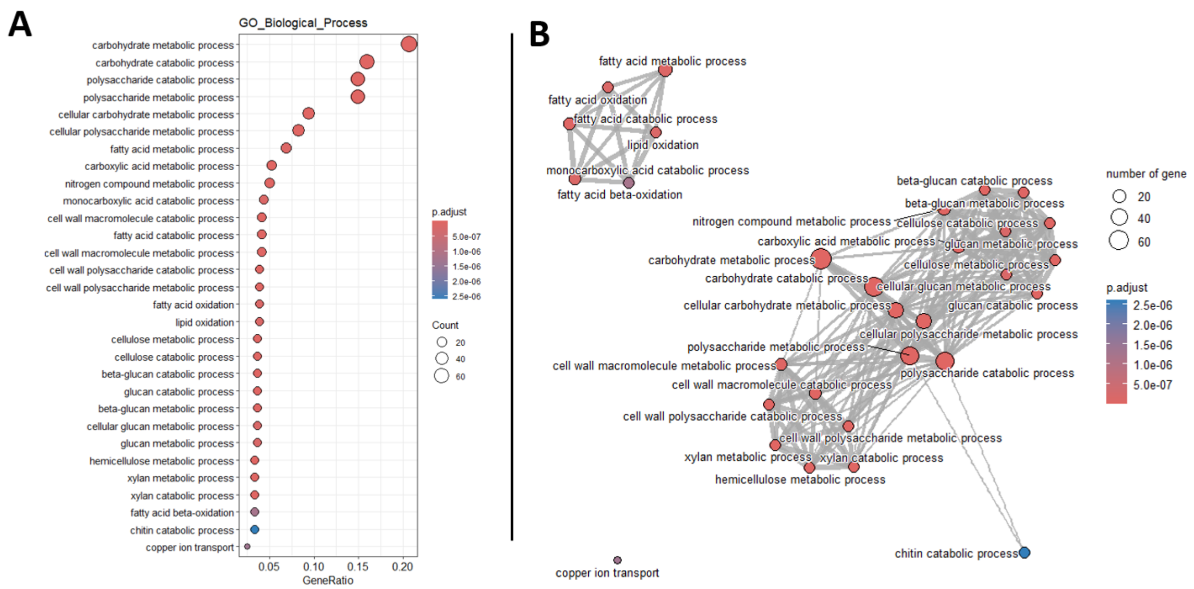


Supplementary Fig. 7. Functional enrichment of genes from expression module 2. Expression modules 2.1, 2.2, 2.3, and 2.4 comprise 451 genes that are upregulated after contact with *A. brassicicola* and *R. solani* in all *T. atroviride* strains, but are more highly expressed in the highly parasitic strains. Dot plot (A) and Enrichment Map (B) of the 30 most significantly enriched GO (Gene Ontology) biological processes.

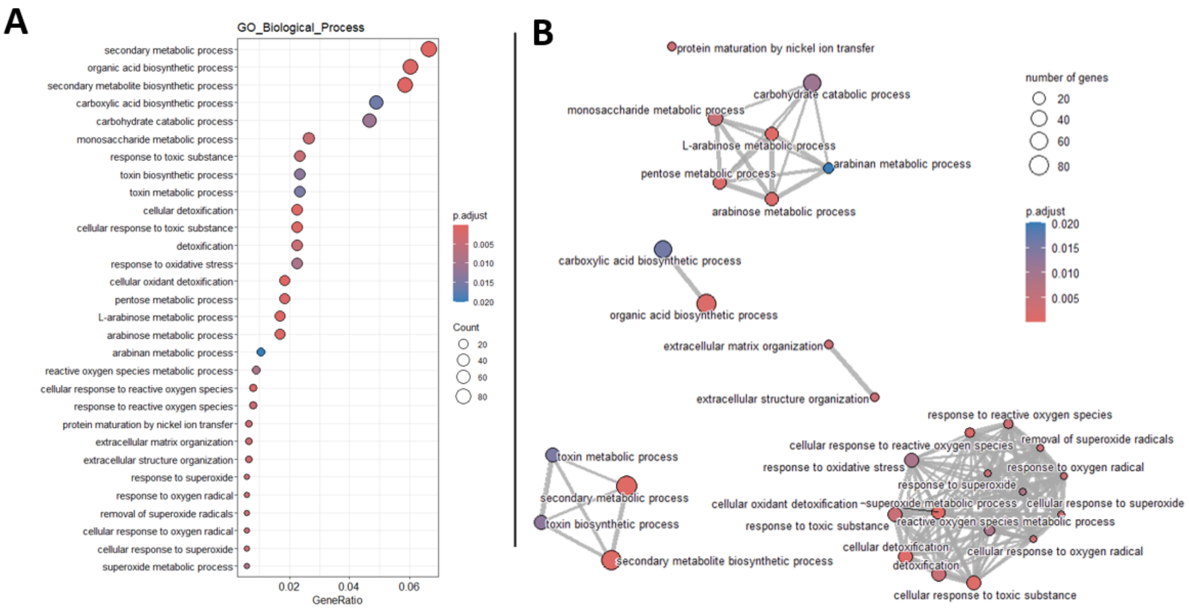


Supplementary Fig. 8. Functional enrichment of genes from expression module 3. Expression modules 3.1, 3.2, 3.3, and 3.4 comprise 1,593 genes that are more highly expressed across all tested conditions in the highly parasitic strains. Dot plot (A) and Enrichment Map (B) of the 30 most significantly enriched GO (Gene Ontology) biological processes.

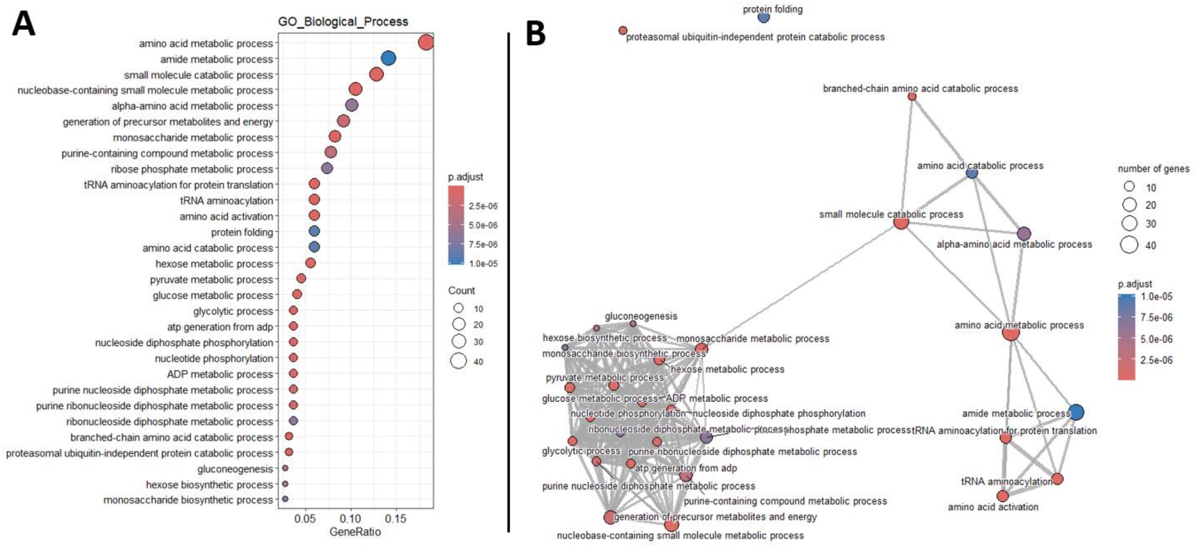


Supplementary Fig. 9. Functional enrichment of genes from expression module 4. Expression modules 4.1, 4.2, and 4.3 comprise 237 genes that are mainly expressed prior to confrontation in the highly parasitic strains and are repressed in response to contact with *A. brassicicola* and *R. solani*. These modules show an expression pattern opposite to that of modules 2. Dot plot (A) and Enrichment Map (B) of the 30 most significantly enriched GO (Gene Ontology) biological processes.

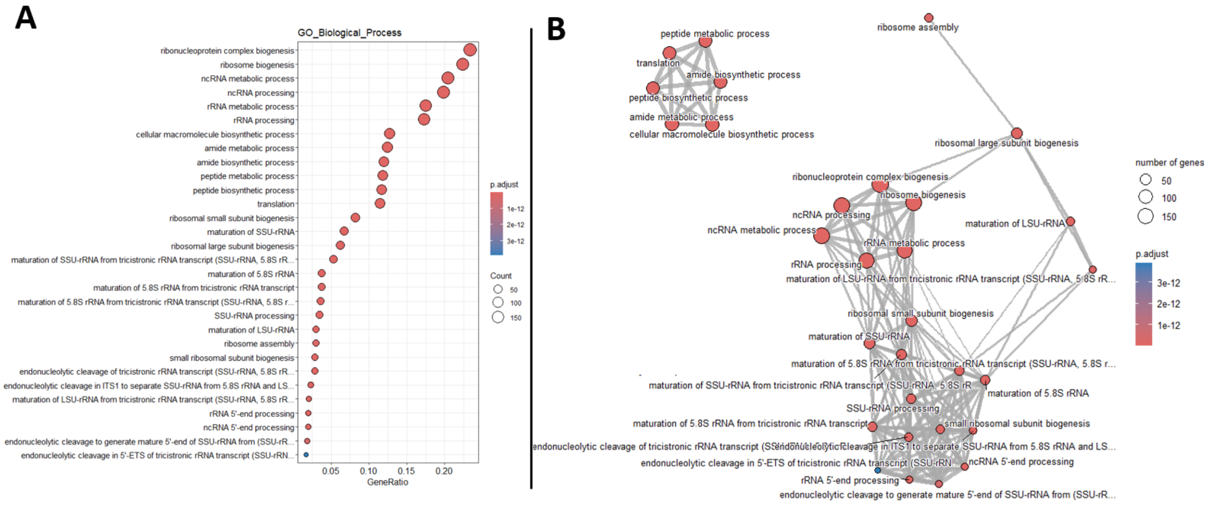


Supplementary Fig. 10. Functional enrichment of genes from expression module 5. Expression modules 5.1, 5.2, 5.3 and 5.4 comprise 862 genes that are more highly expressed before confrontation than after across all strains. Dot plot (A) and Enrichment Map (B) of the 30 most significantly enriched GO (Gene Ontology) biological processes.

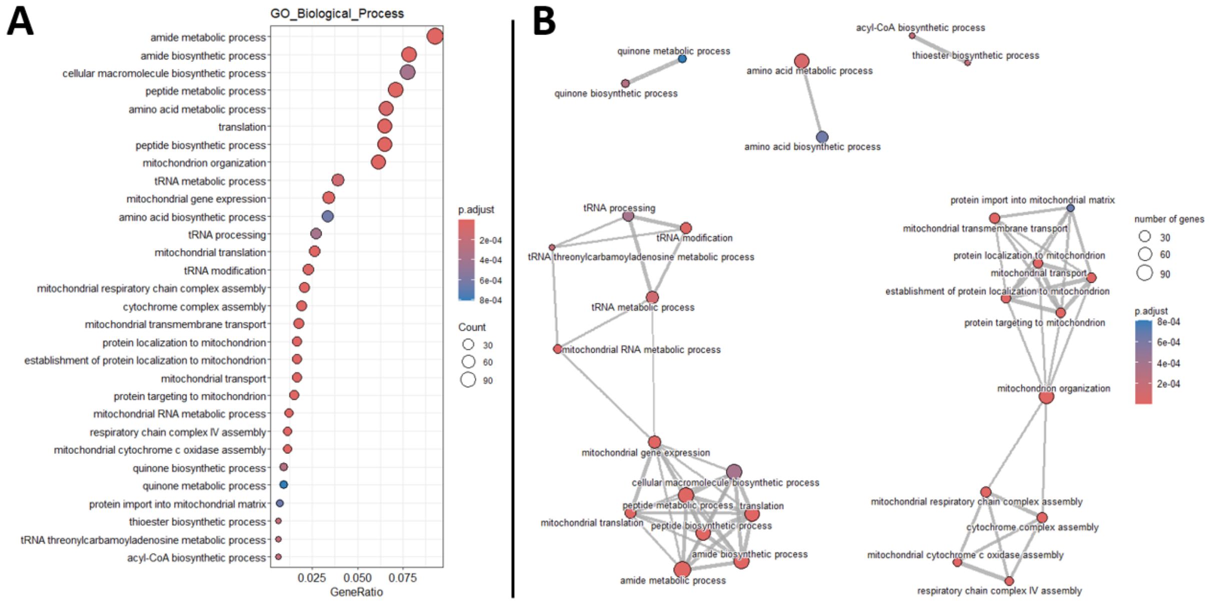


Supplementary Fig. 11. Functional enrichment of genes from expression module 6. Expression modules 6.1, 6.2, 6.3, 6.4, 6.5, and 6.6 comprise 1,568 genes that show lower expression in highly parasitic strains relative to weakly parasitic strains, with minor differences between pre- and post-contact conditions. Dot plot (A) and Enrichment Map (B) of the 30 most significantly enriched GO (Gene Ontology) biological processes.

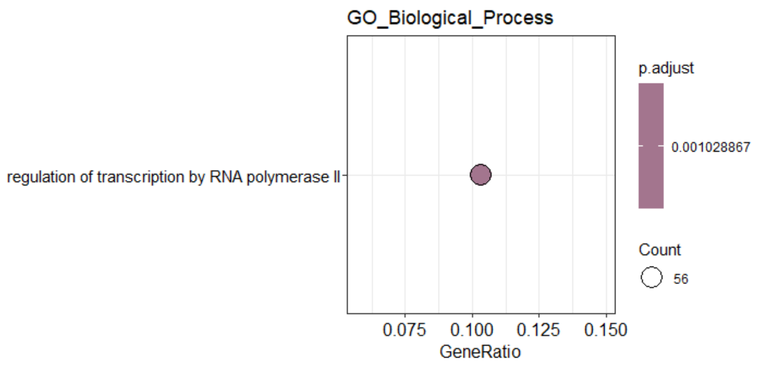


Supplementary Fig. 12. Functional enrichment of genes from expression module 7. Expression modules 7.1, 7.2, 7.3, 7.4, and 7.5 comprise 688 genes that are more variable, with overall lower expression in MMS1295 and N1508 compared to the other strains. This module primarily reflects phylogenetic differences among the six strains. Dot plot of the 30 most significantly enriched GO (Gene Ontology) biological processes.

| Substrate | GH family | Gene count per genome: Weakly parasitic I1237 | Weakly parasitic P3041 | Highly parasitic strains P3080 | Highly parasitic P3116 | Highly parasitic MMS1295 | Highly parasitic N1508 | Genes overexpressed in response to pathogen contact — Up-regulated after contact vs before in WP | HP | or both strains | Differentially expressed genes between WP and HP strains — After contact, higher expression by WP | vs HP strains |
|---|---|---|---|---|---|---|---|---|---|---|---|---|
| α-glucans | GH13 | 5 | 5 | 5 | 5 | 5 | 5 | 1 | 2 | | 1 | 3 |
| | GH132 | 2 | 2 | 2 | 2 | 2 | 2 | | | | | 1 |
| | GH15 | 3 | 3 | 3 | 3 | 3 | 3 | 1 | | | 2 | 1 |
| | GH31 | 7 | 7 | 7 | 7 | 7 | 7 | | | 3 | 3 | 2 |
| | GH32 | 1 | 1 | 1 | 1 | 1 | 1 | | | | | |
| | GH37 | 2 | 2 | 2 | 2 | 2 | 2 | 1 | | | 1 | 1 |
| | GH63 | 1 | 1 | 1 | 1 | 1 | 1 | | | | | |
| | **GH65** | 2 | 2 | 2 | 2 | 2 | 2 | 1 | | 1 | 1 | 1 |
| | **GH71** | 4 | 4 | 4 | 4 | 4 | 4 | | 2 | 2 | | 4 |
| α-mannan | GH125 | 2 | 2 | 2 | 2 | 2 | 2 | 1 | | | 1 | |
| | GH38 | 1 | 1 | 1 | 1 | 1 | 1 | | | | | |
| | GH47 | 8 | 8 | 8 | 8 | 8 | 8 | | | 3 | | 3 |
| | GH76 | 9 | 9 | 9 | 9 | 9 | 9 | 2 | 3 | 3 | 5 | 1 |
| | **GH92** | 8 | 8 | 8 | 8 | 8 | 8 | 3 | | 4 | 1 | 4 |
| Aminoglycan | GH114 | 1 | 1 | 1 | 1 | 1 | 1 | | 1 | | | |
| | GH117 | 1 | 1 | 1 | 1 | 1 | 1 | | | 1 | | |
| | GH89 | 1 | 1 | 1 | 1 | 1 | 1 | | 1 | | 1 | |
| β-glucans | GH128 | 5 | 5 | 5 | 5 | 5 | 5 | | 1 | 2 | | 4 |
| | GH16 | 17 | 17 | 17 | 17 | 17 | 17 | | 7 | 5 | 7 | 4 |
| | GH17 | 4 | 4 | 4 | 4 | 4 | 4 | | 2 | | | 2 |
| | **GH30_2** | 2 | 2 | 2 | 2 | 2 | 2 | | | 2 | | 2 |
| | GH5_gluc | 2 | 2 | 2 | 2 | 2 | 2 | | 1 | | | 1 |
| | **GH55** | 7 | 7 | 7 | 7 | 7 | 7 | | 3 | 3 | | 6 |
| | **GH64** | 3 | 3 | 3 | 3 | 3 | 3 | 1 | | 2 | 1 | 1 |
| | GH72 | 5 | 5 | 5 | 5 | 5 | 5 | | 1 | 1 | | 4 |
| | **GH81** | 2 | 2 | 2 | 2 | 2 | 2 | | | 1 | 1 | 1 |
| β-mannan<br>α-galactosides<br>α-arabinofuranosides<br>β-xylosides<br>α-glucuronosides | GH105 | 2 | 2 | 2 | 2 | 2 | 2 | 1 | | 1 | 1 | 1 |
| | GH115 | 1 | 1 | 1 | 1 | 1 | 1 | | | | | |
| | **GH27** | 9 | 9 | 9 | 9 | 9 | 9 | 2 | | 7 | 2 | 2 |
| | GH30_3 | 1 | 1 | 1 | 1 | 1 | 1 | | 1 | | | 1 |
| | GH36 | 2 | 2 | 2 | 2 | 2 | 2 | 1 | | | 1 | |
| | GH43 | 6 | 6 | 6 | 6 | 5 | 5 | 2 | | 3 | 2 | 2 |
| | GH5_mann | 2 | 2 | 2 | 2 | 2 | 2 | | 1 | 1 | | 2 |
| | GH51 | 1 | 1 | 1 | 1 | 1 | 1 | | | 1 | | |
| | **GH54** | 2 | 2 | 2 | 2 | 2 | 2 | | | 2 | | |
| | GH62 | 2 | 2 | 2 | 2 | 2 | 2 | | | 2 | 1 | |
| | GH67 | 2 | 2 | 2 | 2 | 2 | 2 | 1 | | 1 | 1 | |
| | GH93 | 3 | 3 | 3 | 3 | 3 | 3 | | | 3 | | 2 |
| Cellulose | GH1 | 4 | 4 | 4 | 4 | 4 | 4 | 2 | 1 | 1 | | 4 |
| | GH12 | 3 | 3 | 3 | 3 | 3 | 3 | | 1 | 2 | | 1 |
| | GH3_1 | 12 | 12 | 12 | 12 | 12 | 12 | 2 | 3 | 6 | 2 | 7 |
| | GH45 | 1 | 1 | 1 | 1 | 1 | 1 | | | | | |
| | GH5_cell | 7 | 7 | 7 | 7 | 7 | 7 | 1 | | 4 | | 3 |
| | GH6 | 1 | 1 | 1 | 1 | 1 | 1 | | | 1 | | |
| | GH7 | 2 | 2 | 2 | 2 | 2 | 2 | | | 2 | | |
| Chitin<br>Chitosan | GH18_A | 7 | 7 | 8 | 8 | 7 | 7 | 2 | | 2 | 1 | 3 |
| | **GH18_B** | 13 | 13 | 13 | 13 | 13 | 13 | | 2 | 9 | 3 | 6 |
| | GH18_C | 8 | 8 | 8 | 8 | 11 | 11 | 3 | 1 | | 3 | 3 |
| | GH20 | 3 | 3 | 3 | 3 | 3 | 3 | | 1 | 2 | 1 | |
| | GH3_3 | 2 | 2 | 2 | 2 | 2 | 2 | 1 | | 1 | | |
| | **GH75** | 5 | 5 | 5 | 5 | 5 | 5 | | 1 | 4 | | 2 |
| Intracellular glucosides | GH2 | 10 | 10 | 10 | 10 | 10 | 10 | 5 | | 4 | 2 | 3 |

| | | | | | | | | | |
|---|---|---|---|---|---|---|---|---|---|
| Pectin α-fucosides | GH106 | 1 | 1 | 1 | 1 | 1 | 1 | 1 | |
| | GH127 | 1 | 1 | 1 | 1 | 1 | 1 | 1 | |
| | GH142 | 1 | 1 | 1 | 1 | 1 | 1 | 1 | 1 |
| | GH154 | 2 | 2 | 2 | 2 | 2 | 2 | 1 | 1 |
| | GH28 | 6 | 6 | 6 | 6 | 6 | 6 | 3, 2 | 2, 1 |
| | GH35 | 1 | 1 | 1 | 1 | 1 | 1 | 1 | 1 |
| | GH78 | 3 | 3 | 3 | 3 | 3 | 3 | 2, 1 | 2 |
| | **GH79** | 4 | 4 | 4 | 4 | 4 | 4 | 3, 1 | 2 |
| | GH88 | 2 | 2 | 2 | 2 | 2 | 2 | 2 | 2 |
| | **GH95** | 4 | 4 | 4 | 4 | 4 | 4 | 1, 2 | 2 |
| Peptido-glucan Iduronid | GH152 | 1 | 1 | 1 | 1 | 1 | 0 | | |
| | GH23 | 1 | 1 | 1 | 1 | 1 | 1 | 1 | 1 |
| | GH25 | 1 | 1 | 1 | 1 | 1 | 1 | | 1 |
| | GH39 | 2 | 2 | 2 | 2 | 2 | 2 | 1, 1 | |
| Xylan Xyloglucan | GH10 | 1 | 1 | 1 | 2 | 2 | 2 | 1 | |
| | GH11 | 4 | 4 | 4 | 4 | 4 | 4 | 4 | 1, 2 |
| | GH3_2 | 1 | 1 | 1 | 2 | 1 | 1 | 1 | 1 |
| | **GH30_1** | 2 | 2 | 2 | 2 | 2 | 2 | 2 | |
| | GH74 | 1 | 1 | 1 | 1 | 1 | 1 | 1 | |

Supplementary Fig. 13. Number of glycoside hydrolase (GH) in each genome and their expression. The central panel shows the number of genes identified in each of the six *Trichoderma atroviride* genomes. On the right, the number of genes overexpressed in post-contact conditions with a pathogen compared to pre-contact conditions is indicated for weakly parasitic (WP), highly parasitic (HP) strains, or both. The far-right panel shows the number of genes more highly expressed by WP or HP strains under post-contact conditions with a pathogen. In grey, non-differentially expressed genes. RNA-seq reads were mapped to the N1508 transcriptome, so gene counts in the expression analysis correspond to N1508 genes. Genes are grouped by GH family, as well as by substrate. GH in bold correspond to those identified as overrepresented in the genomes of mycoparasitic *Trichoderma* compared to saprophytic *Trichoderma* (Gruber and Seidl-Seiboth, 2012).

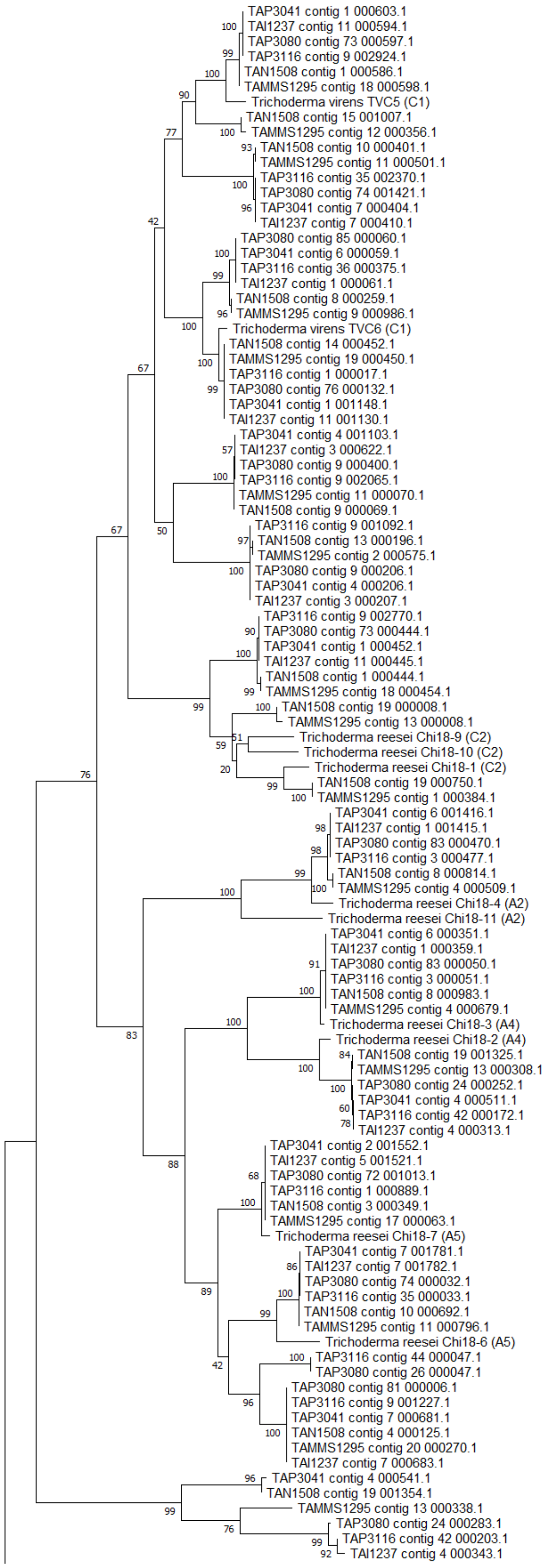
TAP3041 contig 1 000603.1
TAI1237 contig 11 000594.1
TAP3080 contig 73 000597.1
TAP3116 contig 9 002924.1
TAN1508 contig 1 000586.1
TAMMS1295 contig 18 000598.1
Trichoderma virens TVC5 (C1)
TAN1508 contig 15 001007.1
TAMMS1295 contig 12 000356.1
TAN1508 contig 10 000401.1
TAMMS1295 contig 11 000501.1
TAP3116 contig 35 002370.1
TAP3080 contig 74 001421.1
TAP3041 contig 7 000404.1
TAI1237 contig 7 000410.1
TAP3080 contig 85 000060.1
TAP3041 contig 6 000059.1
TAP3116 contig 36 000375.1
TAI1237 contig 1 000061.1
TAN1508 contig 8 000259.1
TAMMS1295 contig 9 000986.1
Trichoderma virens TVC6 (C1)
TAN1508 contig 14 000452.1
TAMMS1295 contig 19 000450.1
TAP3116 contig 1 000017.1
TAP3080 contig 76 000132.1
TAP3041 contig 1 001148.1
TAI1237 contig 11 001130.1
TAP3041 contig 4 001103.1
TAI1237 contig 3 000622.1
TAP3080 contig 9 000400.1
TAP3116 contig 9 002065.1
TAMMS1295 contig 11 000070.1
TAN1508 contig 9 000069.1
TAP3116 contig 9 001092.1
TAN1508 contig 13 000196.1
TAMMS1295 contig 2 000575.1
TAP3080 contig 9 000206.1
TAP3041 contig 4 000206.1
TAI1237 contig 3 000207.1
TAP3116 contig 9 002770.1
TAP3080 contig 73 000444.1
TAP3041 contig 1 000452.1
TAI1237 contig 11 000445.1
TAN1508 contig 1 000444.1
TAMMS1295 contig 18 000454.1
TAN1508 contig 19 000008.1
TAMMS1295 contig 13 000008.1
Trichoderma reesei Chi18-9 (C2)
Trichoderma reesei Chi18-10 (C2)
Trichoderma reesei Chi18-1 (C2)
TAN1508 contig 19 000750.1
TAMMS1295 contig 1 000384.1
TAP3041 contig 6 001416.1
TAI1237 contig 1 001415.1
TAP3080 contig 83 000470.1
TAP3116 contig 3 000477.1
TAN1508 contig 8 000814.1
TAMMS1295 contig 4 000509.1
Trichoderma reesei Chi18-4 (A2)
Trichoderma reesei Chi18-11 (A2)
TAP3041 contig 6 000351.1
TAI1237 contig 1 000359.1
TAP3080 contig 83 000050.1
TAP3116 contig 3 000051.1
TAN1508 contig 8 000983.1
TAMMS1295 contig 4 000679.1
Trichoderma reesei Chi18-3 (A4)
Trichoderma reesei Chi18-2 (A4)
TAN1508 contig 19 001325.1
TAMMS1295 contig 13 000308.1
TAP3080 contig 24 000252.1
TAP3041 contig 4 000511.1
TAP3116 contig 42 000172.1
TAI1237 contig 4 000313.1
TAP3041 contig 2 001552.1
TAI1237 contig 5 001521.1
TAP3080 contig 72 001013.1
TAP3116 contig 1 000889.1
TAN1508 contig 3 000349.1
TAMMS1295 contig 17 000063.1
Trichoderma reesei Chi18-7 (A5)
TAP3041 contig 7 001781.1
TAI1237 contig 7 001782.1
TAP3080 contig 74 000032.1
TAP3116 contig 35 000033.1
TAN1508 contig 10 000692.1
TAMMS1295 contig 11 000796.1
Trichoderma reesei Chi18-6 (A5)
TAP3116 contig 44 000047.1
TAP3080 contig 26 000047.1
TAP3080 contig 81 000006.1
TAP3116 contig 9 001227.1
TAP3041 contig 7 000681.1
TAN1508 contig 4 000125.1
TAMMS1295 contig 20 000270.1
TAI1237 contig 7 000683.1
TAP3041 contig 4 000541.1
TAN1508 contig 19 001354.1
TAMMS1295 contig 13 000338.1
TAP3080 contig 24 000283.1
TAP3116 contig 42 000203.1
TAI1237 contig 4 000343.1

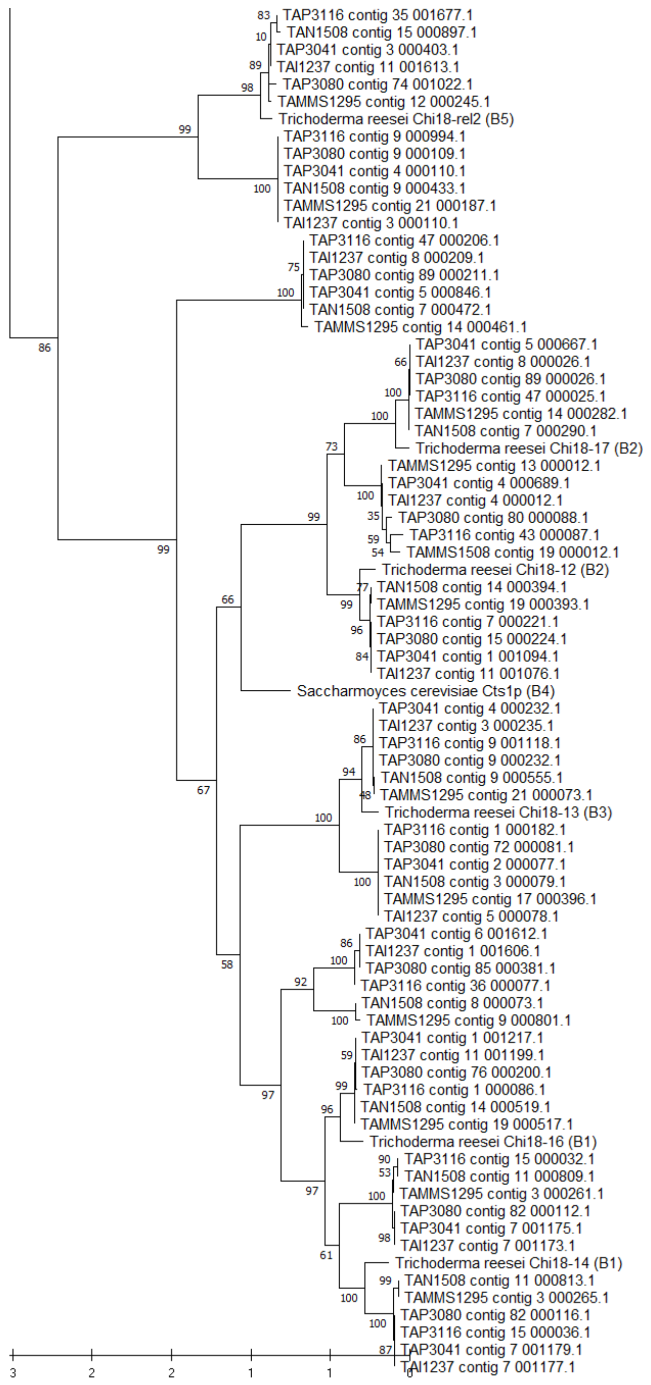


Supplementary Fig. 14. Phylogenetic analysis of GH18 catalytic domains.
The phylogenetic tree was constructed from an amino acid alignment of GH18 catalytic domains. GH18 proteins were first identified using dbCAN3, and catalytic domain sequences were then extracted using PFAM. Reference sequences used by Wang *et al.* (2021) were included for classification, with chitinase group and subgroup indicated in parentheses. Sequences were aligned using MUSCLE, and the tree was inferred in MEGA using the maximum likelihood method with 1,000 bootstrap replicates.

Supplementary Table 2. Number of biosynthetic gene clusters (BGCs) in the genomes of six *Trichoderma atroviride* strains.

BGCs are identified using antiSMASH and classified according to their predicted biosynthetic type. PKS = polyketide synthase; NRPS = nonribosomal peptide synthetase.

| Biosynthetic Gene Cluster (BGC) | | Number of BGC in genome | | | | | |
|---|---|---|---|---|---|---|---|
| | | Weakly parasitic | | Highly parasitic strain | | | |
| | | I1237 | P3041 | P3080 | P3116 | MMS1295 | MMS1508 |
| All BGC | | 48 | 47 | 47 | 47 | 46 | 46 |
| NRPS | NRPS | 8 | 8 | 8 | 7 | 7 | 8 |
| | NRP-metallophore,NRPS | 0 | 0 | 0 | 1 | 1 | 0 |
| | NRPS-like | 5 | 5 | 5 | 5 | 5 | 5 |
| PKS | T1PKS | 11 | 11 | 11 | 11 | 11 | 11 |
| NRPS-PKS | NRPS,NRPS-like,T1PKS | 2 | 2 | 2 | 2 | 2 | 2 |
| | NRPS,T1PKS | 2 | 2 | 2 | 2 | 2 | 2 |
| Isocyanide | isocyanide-nrp | 2 | 2 | 2 | 2 | 2 | 2 |
| | isocyanide | 2 | 2 | 2 | 2 | 1 | 1 |
| | isocyanide,isocyanide-nrp,terpene | 1 | 1 | 0 | 1 | 0 | 0 |
| | isocyanide,isocyanide-nrp | 0 | 0 | 1 | 0 | 1 | 1 |
| Terpene | terpene | 12 | 11 | 11 | 11 | 11 | 11 |
| | terpene-precursor | 2 | 2 | 2 | 2 | 2 | 2 |
| | NRPS,T1PKS,terpene | 1 | 1 | 1 | 1 | 1 | 1 |

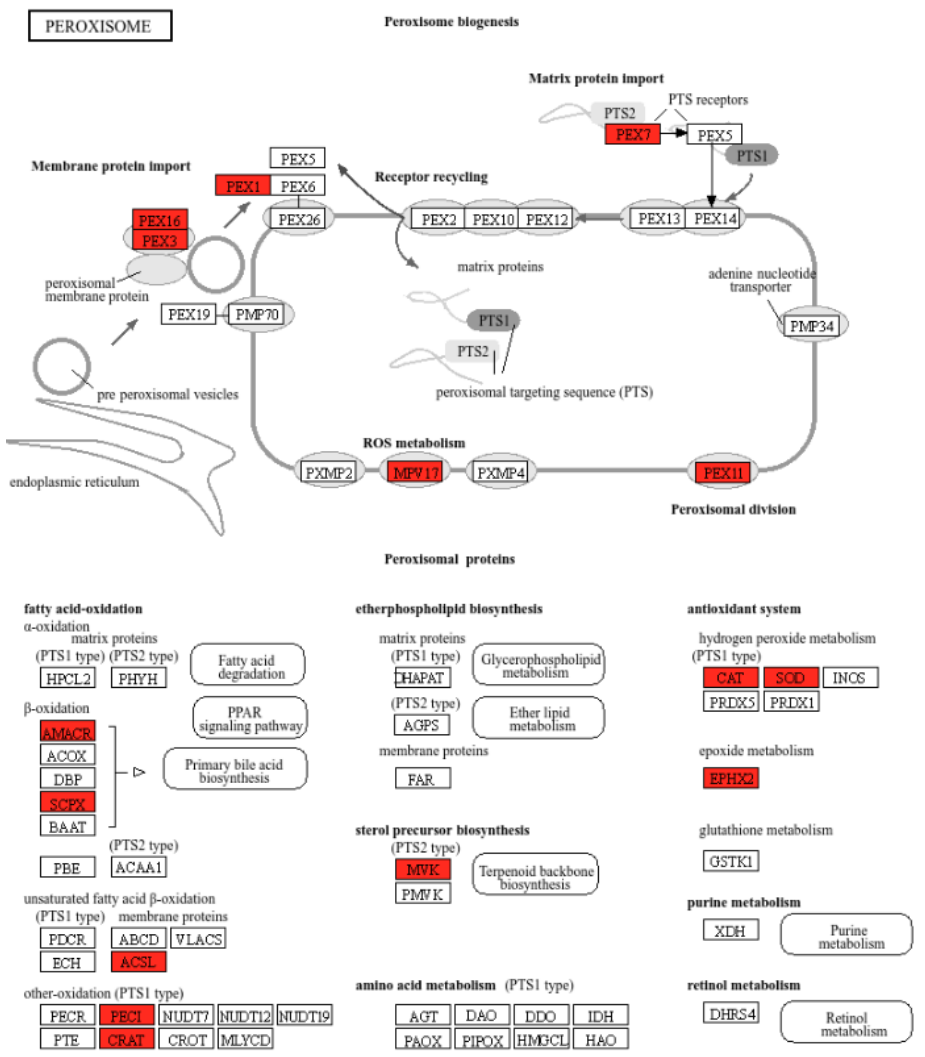


Supplementary Fig. 15. Genes more highly expressed in HP strains in relation to peroxisomes. KEGG map “Peroxisome” showing, in red, genes significantly more highly expressed in highly parasitic strains compared to weakly parasitic strains under post-contact conditions with a pathogen. The upper part shows genes associated with peroxisome biogenesis, and the lower part shows genes associated with the different peroxisomal functions.

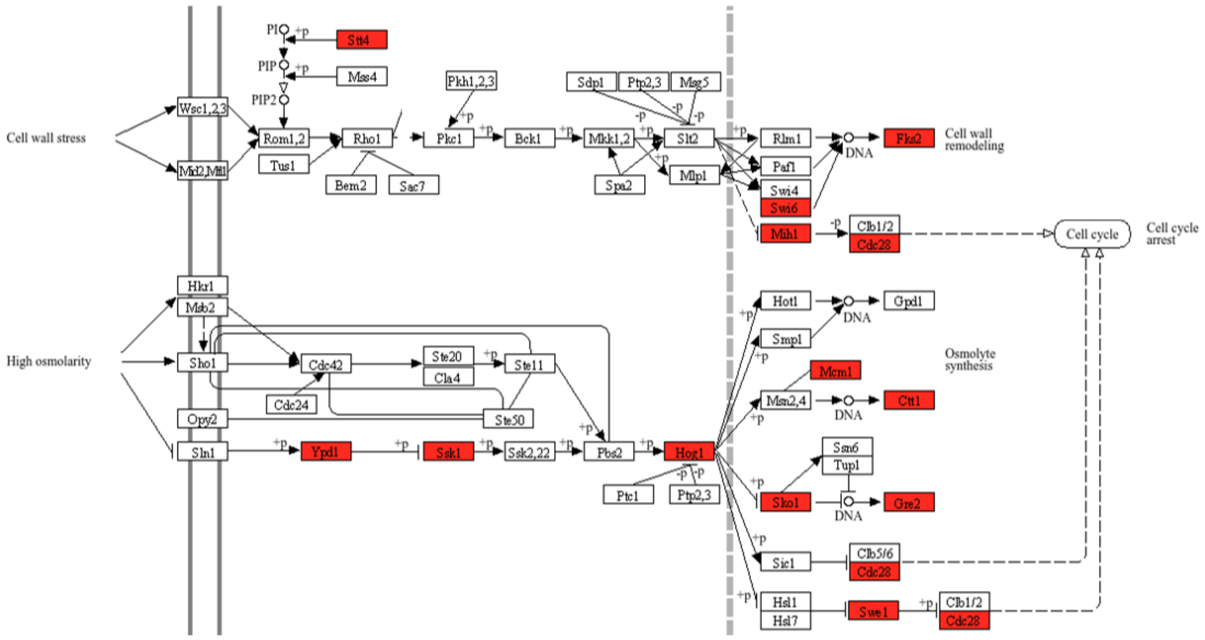


Supplementary Fig. 16. Genes more highly expressed in HP strains in stress response pathways. KEGG map “MAPK signaling pathway – yeast” showing, in red, genes significantly more highly expressed in highly parasitic strains compared to weakly parasitic strains under post-contact conditions with a pathogen. The upper part represents the CWI (Cell Wall Integrity) pathway, and the lower part represents the HOG (High Osmolarity Glycerol) pathway.
